# A systematic survey of PRMT interactomes reveals key roles of arginine methylation in the global control of RNA splicing and translation

**DOI:** 10.1101/746529

**Authors:** Huan-Huan Wei, Xiao-Juan Fan, Yue Hu, Xiao-Xu Tian, Meng Guo, Zhao-Yuan Fang, Ping Wu, Shuai-Xin Gao, Chao Peng, Yun Yang, Zefeng Wang

**Affiliations:** CAS Key Laboratory of Computational Biology, CAS-MPG Partner Institute for Computational Biology, Shanghai Institute of Nutrition and Health, CAS Center for Excellence in Molecular Cell Science, Shanghai Institutes for Biological Sciences, University of Chinese Academy of Sciences, Chinese Academy of Sciences, Shanghai 200031, China; National Facility for Protein Science in Shanghai, ZhangJiang Lab, Shanghai Advanced Research Institute, Chinese Academy of Sciences, Shanghai 201210, China; XiJing hospital of Digestive Diseases, Fourth Military Medical University, XiAn, ShanXi 710000, China

**Keywords:** Post-translational modification (PTM), Protein arginine methyltransferase (PRMT), RNA-binding protein (RBP), alternative splicing, mRNA translation, ribosomal proteins

## Abstract

Thousands of proteins undergo arginine methylation, a widespread post-translational modification catalyzed by various protein arginine methyltransferases (PRMTs). However, a full picture of the catalytic network for each PRMT is lacking and the global understanding of their biological roles remains limited. Here we systematically identified interacting proteins for all human PRMTs and demonstrated that they are functionally important for mRNA splicing and translation. We showed that the interactomes of human PRMTs are significantly overlapped with the known methylarginine containing proteins, and different PRMTs are functionally complementary with a high degree of overlap in their substrates and high similarities between their putative methylation motifs. Importantly, arginine methylation is significantly enriched in RNA binding proteins involved in regulating RNA splicing and translation, and inhibition of PRMTs leads to global alteration of alternative splicing and suppression of translation. In particular, ribosomal proteins are pervasively modified with methylarginine, and mutations on their methylation sites suppress ribosome assembly, translation, and eventually cell growth. Collectively, our study provides a novel global view of different PRMT networks and uncovers critical functions of arginine methylation in the regulation of mRNA splicing and translation.

## Introduction

Arginine N-methylation was first discovered in the early 1970s ^1-3^ and later was recognized as a widespread post-translational modification (PTM) in many proteins ^4-9^. It is catalyzed by a class of enzymes known as protein arginine methyltransferases (PRMTs), which covalently link methyl groups to the arginine side chains. Although arginine methylation does not alter the electric charge, it increases amino acid bulkiness and protein hydrophobicity, thus can affect how proteins interact with their partners. This type of PTM has been found to play key roles in various cellular processes, including phase separation, DNA damage repair, transcriptional regulation and RNA metabolism. ^6-13^. As a result, arginine methylation has a profound effect on human diseases such as cancer ^9, 14, 15^ and cardiovascular diseases ^16^.

Nine PRMTs, PRMT1 to PRMT9, have been identified in the human genome (Fig. 1A), which were classified into three types according to the final methylarginine products. Type I PRMTs, including PRMT1, 2, 3, 4, 6, and 8, catalyze the formation of ω-N^G^, N^G^-asymmetric dimethylarginine (aDMA). Type II PRMTs, including PRMT5 and PRMT9, catalyze the formation of ω-N^G^, N^G^-symmetric dimethylarginine (sDMA). PRMT7 is the only type III PRMT and catalyzes ω-N^G^-monomethylarginine (MMA). Methylation of arginine by PRMTs consumes a great deal of cellular energy (12 ATPs for each methyl group added) ^4^ and is found in >10% of all human proteins ^17^, implying an essentail role of arginine methylation in cell growth and preliferation.

**Figure 1.**
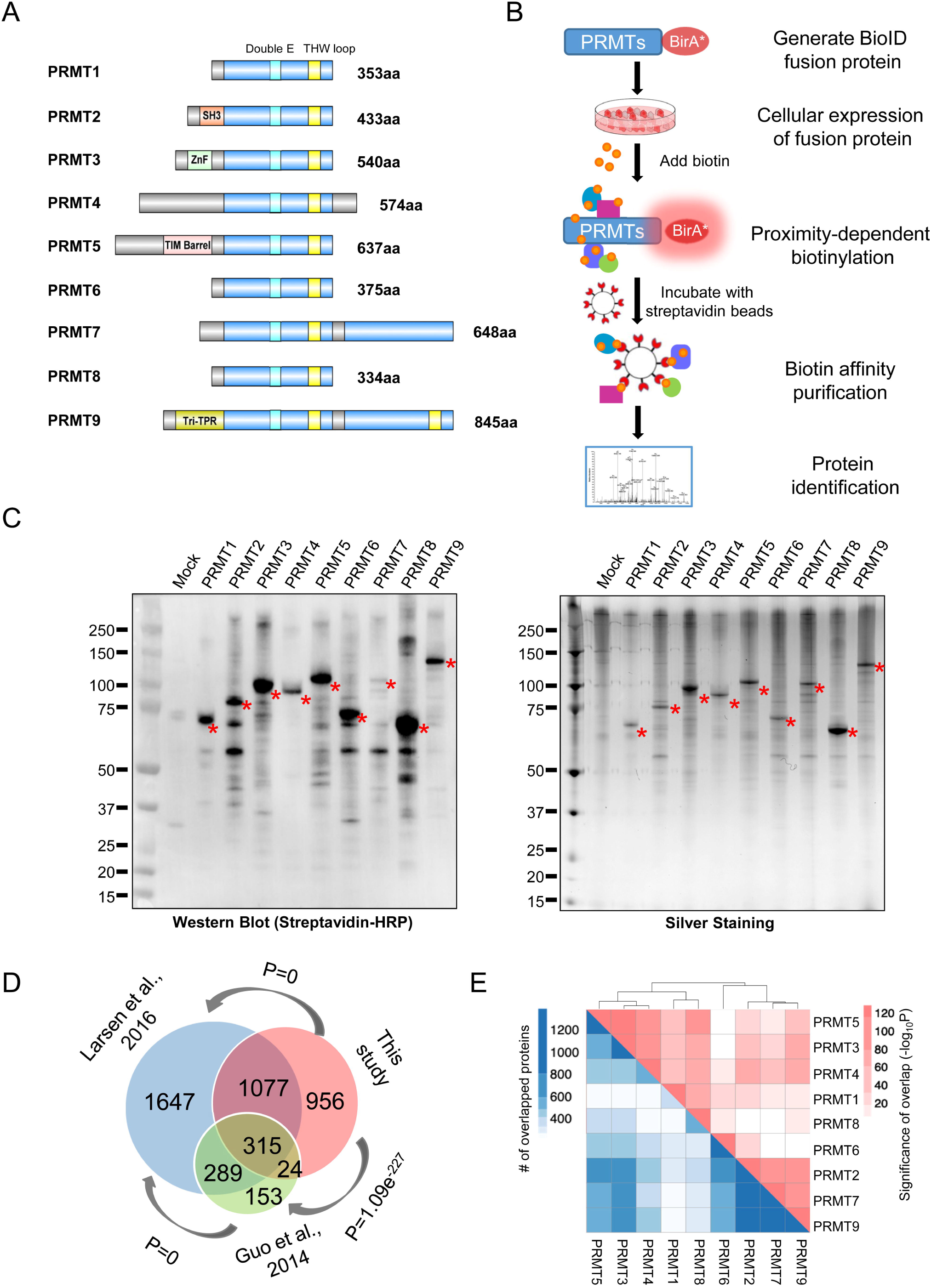
Systematic identification of PRMT interactome. (A) Schematic diagram of PRMT1 to PRMT9. The light blue boxes represent the catalytic domains, the cyan and yellow boxes represent Double E Motifs and THW loop Motifs that are specific to PRMTs, respectively. (B) The workflow for identification of PRMT interacting proteins via BioID. (C) The biotinylated PRMT interacting proteins as detected by western blot using streptavidin-HRP (left) and by silver staining (right). (D) The Venn diagram illustrating the PRMT interactome from this study compared to methylarginine-containing proteins identified in Larsen et al., 2016 and Guo et al., 2014. The correlation is calculated by Fisher’s exact test, with the whole genome as the background. (E) The overlap of the interactome among different PRMTs. Fishers’ exact test is used to calculate the p-value of the overlap. The PRMT interactomes are clustered by overlap significance (shown in red) and the numbers of overlapped protein are indicated in blue.

The biological functions of PRMTs are largely determined by its substrates and regulating partners, and therefore identifying the full scope of the interactors for each PRMT will greatly improve our understanding of the function of arginine methylation. The substrates of several individual PRMTs (e.g., PRMT4 and PRMT5) have been determined using various approaches, however, the substrates or interactors of other PRMTs remain largely unknown. On the other hand, thousands of human proteins have been identified to undergo arginine methylation using mass spectrometry combined with specific methylarginine antibodies ^17, 18^, and thus efforts to connect these arginine metylated proteins with their respective PRMTs are highly valuable to further understand their regulation and function.

In this study, we systematically identified interactome of each PRMT using BioID that allows identification of transient protein-protein interactions ^19-21^, and further determined the substrate specificity and consensus arginine methylation motifs of each PRMT. Our results revealed a high degree of overlap in substrate specificity of different PRMTs, suggesting a possible functional complementation. Remarkably, PRMT interactors are significantly enriched for RNA binding proteins involved in mRNA splicing and translation, and inhibition of PRMTs leads to global alteration of alternative splicing and reduction of mRNA translation. Particularly, mutations on methylation site of ribosomal proteins inhibited ribosome assembly and cell growth. Our study therefore systematically links individual PRMT to its arginine methylation events, and highlights the importance of PRMTs in regulation of RNA metabolism.

## Results

### Identification of the interacting proteins of each PRMT

As protein modification enzymes, PRMTs dynamically interact with their protein substrates, making it difficult to identify these interacting partners. To systematically characterize interactome of different PRMTs *in vivo*, we applied the highly sensitive BioID technology to label interacting proteins with biotin. Based on previous reports that N-terminus of PRMT is responsible for substrate recognition ^22-25^, we fused a promiscuous biotin ligase BirA* (BirA^R118G^) to the C-terminus of each PRMT and expressed the fusion proteins in HEK293T cells. Using cell fractionation followed by western blotting, we confirmed that the fusion proteins have a similar pattern of cytoplasmic *vs.* nuclear localization compared to endogenous PRMTs (Fig. S1). After transient expression of each fusion protein, we purified the total biotinylated proteins with streptavidin beads using a stringent washing condition (2% SDS), and subsequently conducted mass spectrometry with label-free quantitative (LFQ) (Fig. 1B, and experimental procedures). We performed the transfection of all PRMT-BirA* fusion proteins and mock control (BirA* only) in three biological replicates, and identified putative interacting proteins of each PRMT that are relatively depleted in the control (see method).

We first confirmed that the proteins purified from the cells transfected with PRMT-BirA* are indeed heavily biotinylated as compared to the BirA* control using western blotting (Fig. 1C, left), with the PRMT themselves being the most heavily biotinylated proteins (Fig. 1C, right). The LFQ intensities of all detected proteins (as calculated by label free quantification) are highly correlated between different replicates of the same PRMT (Fig. S2A), suggesting that the identification by this procedure is fairly reliable.

In total, we have identified 2372 candidate proteins bound by at least one of the nine PRMTs (Table S1 and Fig. S2B). The identified proteins include many known partners required for PRMT activities, such as the MEP50 (WDR77), RIOK and ICLn, which form an active methyltransferase complex with PRMT5 ^26, 27^. In addition, many of the identified PRMT interactors significantly overlapped with the proteins identified in the earlier proteomic studies using antibodies against methylarginine-containing oligopeptides ^17, 18^ (Fig. 1D), indicating the BioID technology is sensitive enough to identify their substrates that may transiently interact with the enzymes. These proteins include many of the known PRMT substrates including several histones and hnRNPs (table S1). Further analysis of the amino acid composition further showed that the newly identified PRMT interactors contain a higher fraction of arginine as compared to all human proteins, again suggesting an enrichment of PRMT substrates (Fig. S2C).

In addition to the methylation substrates, the interactome of PRMTs may also include proteins that regulate PRMT functions, which could be missed in immunoprecipitation with methylarginine antibodies. Notably, only 4% of our newly identified PRMT interacting proteins (85 out of 2372 proteins) have been collected in the IntAct online PPI database (https://www.ebi.ac.uk/intact/) (Fig. S2D), indicating that our results significantly expanded the interactome of each PRMT.

### Substrate preference of individual PRMTs

To further determine the substrate preference of the putative substrates for different PRMTs, we compared the newly identified putative substrates for each PRMT. Our results indicated that many proteins are recognized by multiple PRMTs, suggesting a great deal of substrate redundancy for each PRMT (Fig. S2E). For example, 90 proteins could be recognized by all 9 PRMTs as judged by our results, and 961 proteins could be recognized by at least 4 out of 9 PRMTs tested (Fig. S2F). We further examined the overlaps of the interacting proteins between each pair of the PRMTs (Fig. 1E, red, overlap significance; blue, the numbers of overlapped proteins), and found that the PRMTs can be roughly clustered into two groups based on the similarity of their interactomes, with one group containing PRMT2, PRMT6, PRMT7 and PRMT9, whereas another group containing PRMT1, PRMT3, PRMT4, PRMT5 and PRMT8.

### Identification and validation of consensus motifs for arginine methylation

Previous studies have reported that the glycine and arginine rich (GR-rich) motifs are preferably targeted for methylation by many PRMTs (including PRMT1, PRMT3, PRMT5, PRMT6, and PRMT8) ^9, 28, 29^. However, additional consensus motifs such as proline/glycine/methionine rich (PGM-rich) or RxR motifs were also found to be enriched near the methylarginine sites by mass spectrometry ^30, 31^, suggesting that other sequences beside GR-rich motifs may also be recognized as arginine methylation sites and that individual PRMTs may have different preferences of their substrate.

To further determine the substrate preference for different enzymes, we analyzed the newly identified putative PRMT substrates by measuring the statistic enrichment of the sequences around the potential methylarginine (Fig. 2A, see methods for details). For each PRMT, the tetrapeptides around the potential methylarginine sites were compared with the arginine-containing tetrapeptides in all proteins from UniProt database to calculate the enrichment Z-scores (Fig. 2A). The enriched tetrapeptides were further clustered into different groups to obtain consensus motifs for arginine methylation by each PRMT. As an example, the clusters and the consensus motifs for PRMT3 substrates were shown in Fig. 2B and the clusters of all PRMTs were shown in supplementary figure S3. We further compared the consensus motifs of all tested PRMTs (Fig. 2C), and found that in addition to the known RGG motifs from the substrates of many PRMTs, several other new consensus motifs like SR-rich, DR-rich, and ER-rich motifs were also be identified in PRMT substrates. These results provided a comprehensive profile for the substrate preference of different PRMTs, suggesting that a diverse range of proteins could be potentially modified by PRMTs at different consensus motifs.

**Figure 2.**
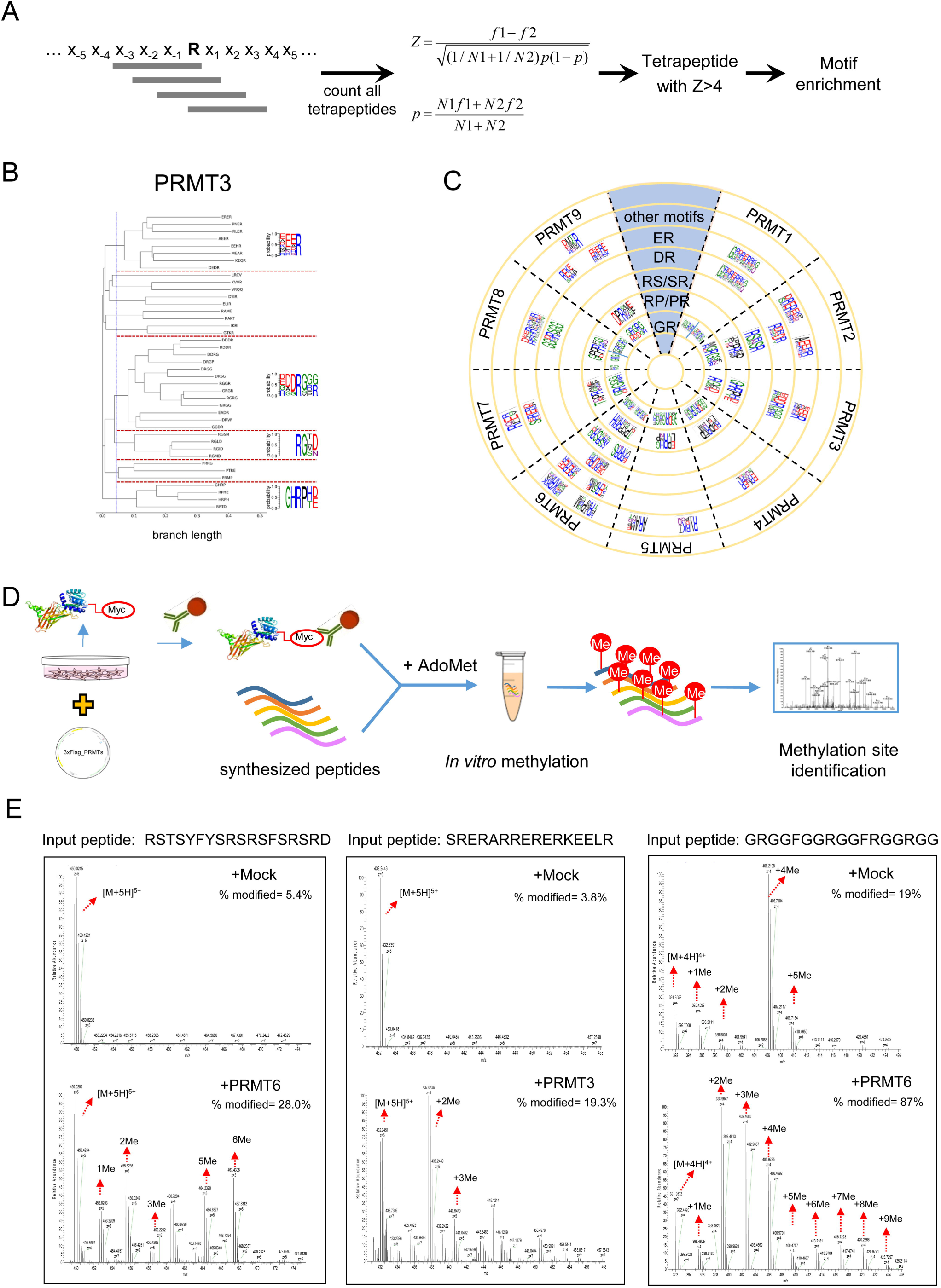
Identification and validation of consensus motifs for arginine methylation. (A) The schematic diagram for identification of the enriched motif in PRMT interactome. All tetrapeptides around arginine are counted, and the frequencies of each tetrapeptide in PRMT interactomes were compared to the background of all human proteins to identify enriched tetrapeptides (see methods). (B) The clustering of enriched tetrapeptides in PRMT3 interactome is shown as an example. The enriched tetrapeptides are collected as input in clustalw2 (v2.0.9) to generate the phylogenetic tree, and the consensus motifs were listed besides each group. (C) A summary of all enriched motifs found in each PRMT interactome. Similar motifs are placed in the same concentric circle. (D) The experimental workflow of *in vitro* methylation and identification of methylarginine-containing peptides. R^Me^ and R^2Me^ are set as dynamic modifications with a mass shift of 14.01565 and 28.0313. (E) The arginine methylation pattern of representative peptides from the newly identified R-methylation motifs (SR-rich and ER-rich motifs) and the known GR-rich motif were measured by mass spectrum. In each case, the upper spectrum indicates the negative control without adding enzymes, and the lower spectrum shows the methylarginine signals after *in vitro* methylation. For each peptide, the ratio of methylation was calculated as the sum of the peak areas from the TIC values of the modified peptides divided by the peak area of the total peptides.

In order to validate these newly identified arginine methylation motifs, we selectively synthesized several peptides containing newly identified consensus motifs based on our database to measure methylation of arginine by the cognate PRMTs using *in vitro* methylation reaction (Fig. 2D). The peptides were incubated with purified PRMTs (Fig. S4A and S4B) in the presence of methyl donor S-adenosylmethionine, and resulting samples were analyzed with mass spectrometry. We found that the arginine residues within the SR-rich, ER-rich, and GR-rich motifs can be robustly methylated by cognate PRMTs, with both methylation and dimethylation being detected (Fig. 2E), indicating that these newly identified consensus motifs can indeed be methylated at the arginine sites.

### Potential functions of PRMT substrates

To examine the functional consequence of arginine methylation, we inferred the potential functions of the newly identified PRMT substrates using gene ontology (GO) analyses (https://david.ncifcrf.gov/) ^32, 33^. In order to increase the specificity of our analysis and reduce the statistic noises from the large number of potential substrates, we first focused on the proteins that were identified in both our dataset and from earlier reports of methylarginine-containing proteins ^17, 18^. We found that these proteins were significantly enriched for biological processes involving RNA metabolisms, such as mRNA splicing, translation initiation and mRNA export (Fig. 3A, left). Consistently, these proteins are also enriched for RNA binding domains such as RNA recognition motif (RRM), DEAD/DEAH RNA helicase, KH domain (Fig. S5). The significant involvement of PRMT substrates in RNA metabolism supported the previous reports that many proteins with methylarginine modification participate in RNA processing ^17, 18^. In addition, other PRMT interactors that do not overlap with previously reported methylarginine-containing proteins are likely to be the regulator of PRMTs rather than their substrates (e.g. WDR77). Interestingly, these proteins are enriched for functions related to cell division/cell cycle and NFκB signaling (Fig. 3A, right). Consistently, these proteins are more enriched for domains involved in protein-protein interactions (e.g. Armadillo-type fold) and the signal transduction (e.g. Ser/Thr protein kinase, TPR-like repeats) (Fig. S5).

**Figure 3.**
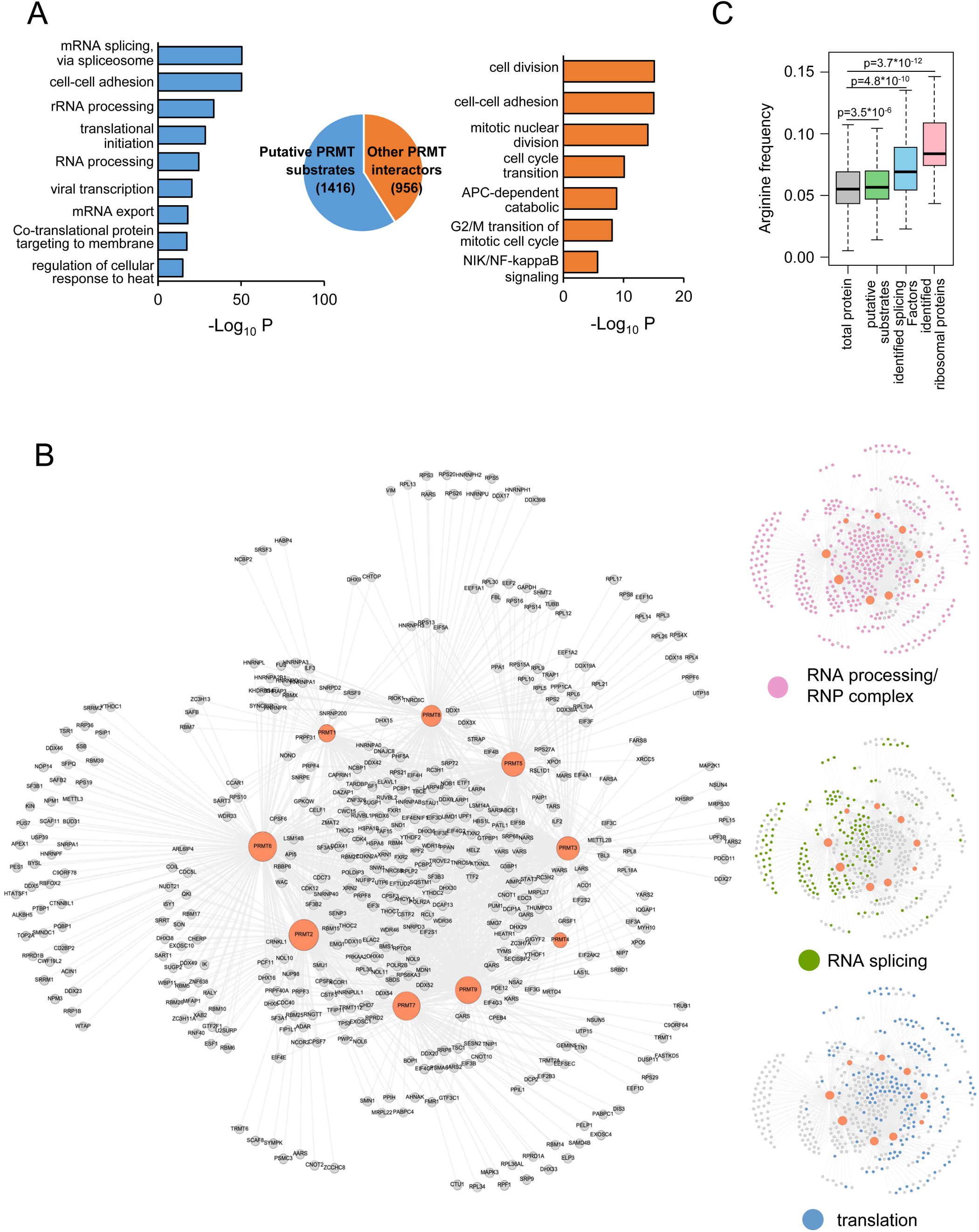
Arginine methylation is highly involved in RNA splicing and translation. (A) Gene Ontology analysis (by DAVID) of the 1416 putative PRMT substrates detected in both this study and previously identified methylarginine-containing proteins (Putative PRMT substrates, blue bars), as well as other PRMT interacting proteins identified in only this study (orange bar). (B) Connectivity map of nine PRMTs (brown circles) and their putative substrates (Left). The putative substrates were also classified and colored according to their major functions (Right). (C) Arginine frequency of all annotated human proteins (n= 42119), identified putative PRMT substrates (n=1416), identified splicing factors with methyl-arginine (n=148), and identified ribosomal proteins as PRMT substrates (n=42). Wilcoxon test was used to calculate the *p*-value.

Furthermore, we focused on the candidates that form dense protein-protein interactions (PPI) networks as judged by STRING database. We plotted the putative substrates in connection with each PRMT (Fig. 3B), and again found that most of these proteins primarily involved in RNA processing and formation of RNP complex. These proteins can be roughly divided into two main groups, with one group mainly containing the splicing regulatory factors and the other including proteins in mRNA translation such as the ribosomal proteins (Fig. 3B). In addition, the core ribosomal proteins and the splicing factors identified in our study generally have a significantly higher frequency of arginine in their amino acid composition as compared to all human proteins (Fig. 3C), further supporting the prevalent methylarginine modification observed in these proteins.

### PRMT inhibitions generally altered alternative splicing of RNA

The majority of human genes undergo alternative splicing (AS), which is generally regulated by various RNA-binding proteins (i.e., splicing factors) that recognize regulatory *cis*-elements to promote or suppress the use of adjacent splice sites ^34^. It was previously reported that some PRMTs (e.g., PRMT4 and PRMT5) can affect splicing by modifying selected splicing factors or proteins involved in spliceosome maturation ^30, 35, 36^. Since many proteins involved in splicing regulation were identified as PRMT substrates (Fig. 3B), we examined the impact of PRMT inhibition on alternative RNA splicing. We achieved effective gene silence with shRNAs in six different PRMTs (Fig. S6A) and examined their effect on splicing using RNA-seq (Fig. 4A). For each PRMT we identified the alternative splicing events that are significantly altered in cells with PRMT knockdown compared to the control cells with scramble RNAi (Fig. 4A).

**Figure 4.**
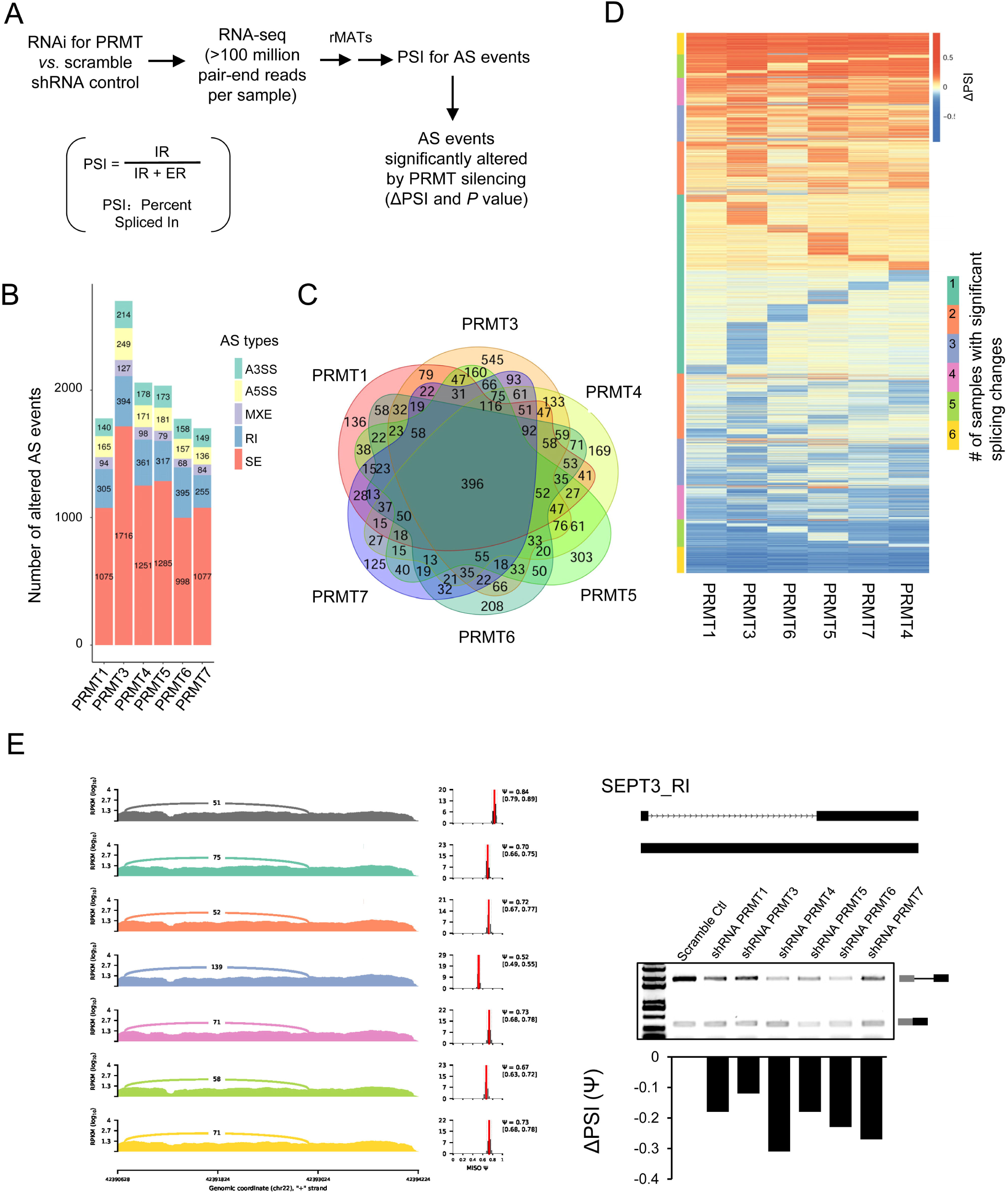
PRMT inhibition leads to global alteration of alternative splicing. (A) The workflow of RNA splicing analysis for PRMT knockdown samples using RNA-seq and analyzed by MISO pipeline to calculate PSI (percent spliced in) values. We used |ΔPSI| >0.1 and read counts>50 as the cutoff to identified significantly altered splicing events. (B) The count of different types of altered splicing events after PRMT knockdown. A3SS, alternative 3′ splice site; A5SS, alternative 5′ splice site; MXE, mutually exclusive exon; RI, retained intron; SE, skipped exon. (C) The intersection of altered AS events upon silencing of different PRMTs. (D) All AS events altered upon silencing of each PRMT were colored according to ΔPSI. The AS events were also clustered by the numbers of PRMT RNAi samples with the significant splicing changes (e.g., the clusters labeled in yellow include AS events affected by RNAi of all the six PRMTs tested). (E) Experimental validation of spicing alteration. Sashimi plot of splicing change in SEPT3 was presented in the left, including the counts of junction read, the PSI value and its confidence interval. Semi-quantitative PCR was shown in the right. Additional examples of altered AS events can be found in fig. S6B

We found that RNAi of different PRMTs led to significant changes of splicing in hundreds of genes harboring various AS types (Fig. 4B). Many of AS events are affected by the inhibition of more than one PRMT (i.e., large overlaps between splicing targets of different PRMTs), suggesting that the arginine methylation of proteins by PRMTs play a general role to regulate alternative splicing (Fig. 4C). Interestingly, the inhibition of different PRMTs generally had similar effects on splicing of specific genes (i.e., the ΔPSI are either positive or negative in most affected genes, Fig. 4D), implying that the methylation of same RNA-binding proteins by different PRMTs produces similar effects on their activities. The splicing changes of selected AS events were further validated using semi-quantitative RT-PCR. For example, the splicing of a retained intron in SEPT3 (neuronal-specific septin-3) gene is promoted by inhibition of all six PRMTs tested (Fig. 4E), and many other genes have undergone alteration of splicing in the same direction (Fig. S6B), supporting the consistent regulation of splicing by different PRMTs.

### Ribosomal proteins are pervasively arginine methylated

Our gene ontology and protein-protein interaction analyses further revealed that proteins involved in mRNA translation are significantly enriched in the newly identified PRMT substrates, including >50% core components of ribosomes (42 out of 80 ribosome proteins) and many canonical translation factors (such as EIF4G1, EIF4B, EIF2A, etc,). In table S2, we listed all the 80 human core ribosome proteins with newly identified interacting PRMTs and the putative methylarginine sites. Our finding is consistent with earlier reports in the late 1970s that both subunits of the ribosome contain methyl arginine as judged by chromatography of short peptides or amino acid residues originated from ribosomal proteins ^37-39^. More recently, several ribosomal proteins were also reported to contain methylarginine, including yeast RPL12 and RPS2 ^40^ and human RPS3 and RPS10 ^41, 42^. The prevalent arginine methylation in core ribosomal proteins we observed suggests a critical role of this type of PTM in protein translation.

To directly test this hypothesis, we performed polysome profiling to isolate different ribosome fractions (the 40S, 60S, 80S, and polysome, Fig. 5A) and evaluated their methylation status by pan-arginine methylation antibodies (Fig. 5B). Our data demonstrated that ribosomal proteins (most having a MW range of 10-50 kD) are pervasively R-methylated in different ribosome profiling fractions (Fig. 5B). Moreover, many ribosomal proteins were detected in the precipitates pulled down by different types of methylarginine antibodies (MMA, aDMA and sDMA antibodies) (Fig. 5C), further supporting our conclusion that ribosome proteins are pervasively methylated at arginine residues. Our observation that most PRMTs were not co-purified with ribosomes (Fig 5A) is consistent with the absence of PRMTs from the known ribo-interactome ^43^. Interestingly, a small fraction of PRMT3 and PRMT5 are co-separated with both ribosomal subunits (40S and, to a lesser extent, 60S) but not with the assembled 80S ribosome and polysomes (Fig. 5A, indicated by asterisks), which is consistent with that PRMT3 can form an active enzyme complex together with RPS2 ^44^.

**Figure 5.**
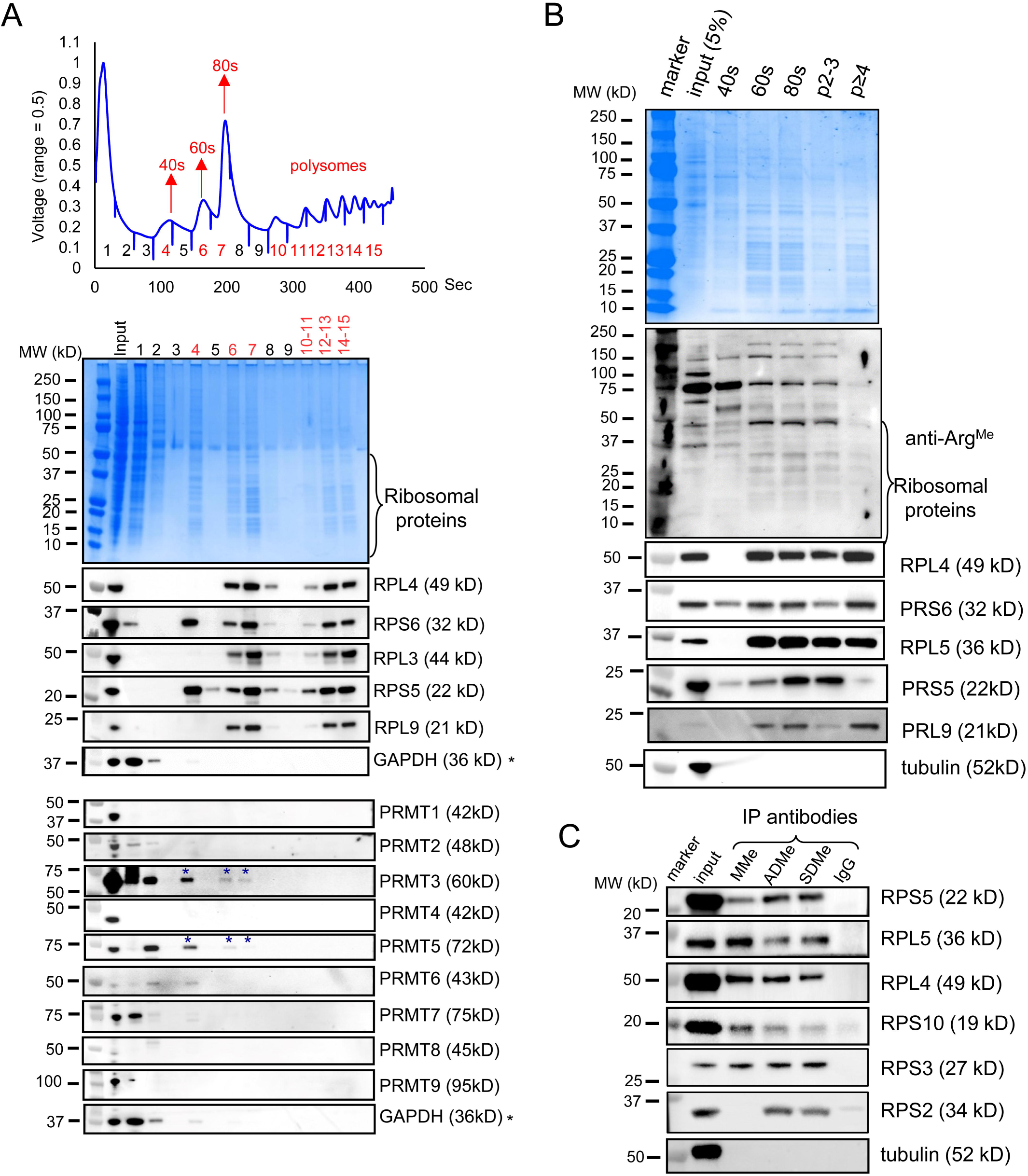
Ribosomal proteins are pervasively methylated. (A) Fractionation of polysomes using HEK293T cell lysis. Each fraction was collected, and proteins in each fraction were precipitated for SDS-PAGE assay. Both coomassie blue staining (top) and western blots (bottom) were used to detect the proteins in each fraction. Accumulation of ribosomal proteins can be observed on the gel (MW between 15-50 kD, Middle). (B) The arginine methylation of ribosomal proteins as detected by combination of pan-methylarginine antibodies that can recognize MMA, aDMA and sDMA. (C) The HEK293T cell lysate were subjected to immunoprecipitation with different methylarginine antibodies, and the selected ribosomal proteins were detected with western blot.

### Arginine methylation is critical for assembly and function of ribosomes

Based on the pervasive arginine methylation of ribosomal proteins and translation factors, we hypothesized that PRMT inhibition may affect translation on a global scale. To test this hypothesis, we used puromycin incorporation assay to test global protein synthesis in three different cell lines after treatment by the PRMT specific inhibitors (Fig. 6A, Fig. S7A and Fig. S7B for cell lines HEK 293T, U2OS and PC9, respectively). We used chemical inhibitors of PRMTs because they could provide a more rapid suppression of ribosomal activity that might otherwise be compensated in cells with stable knockdown of PRMTs.

**Figure 6.**
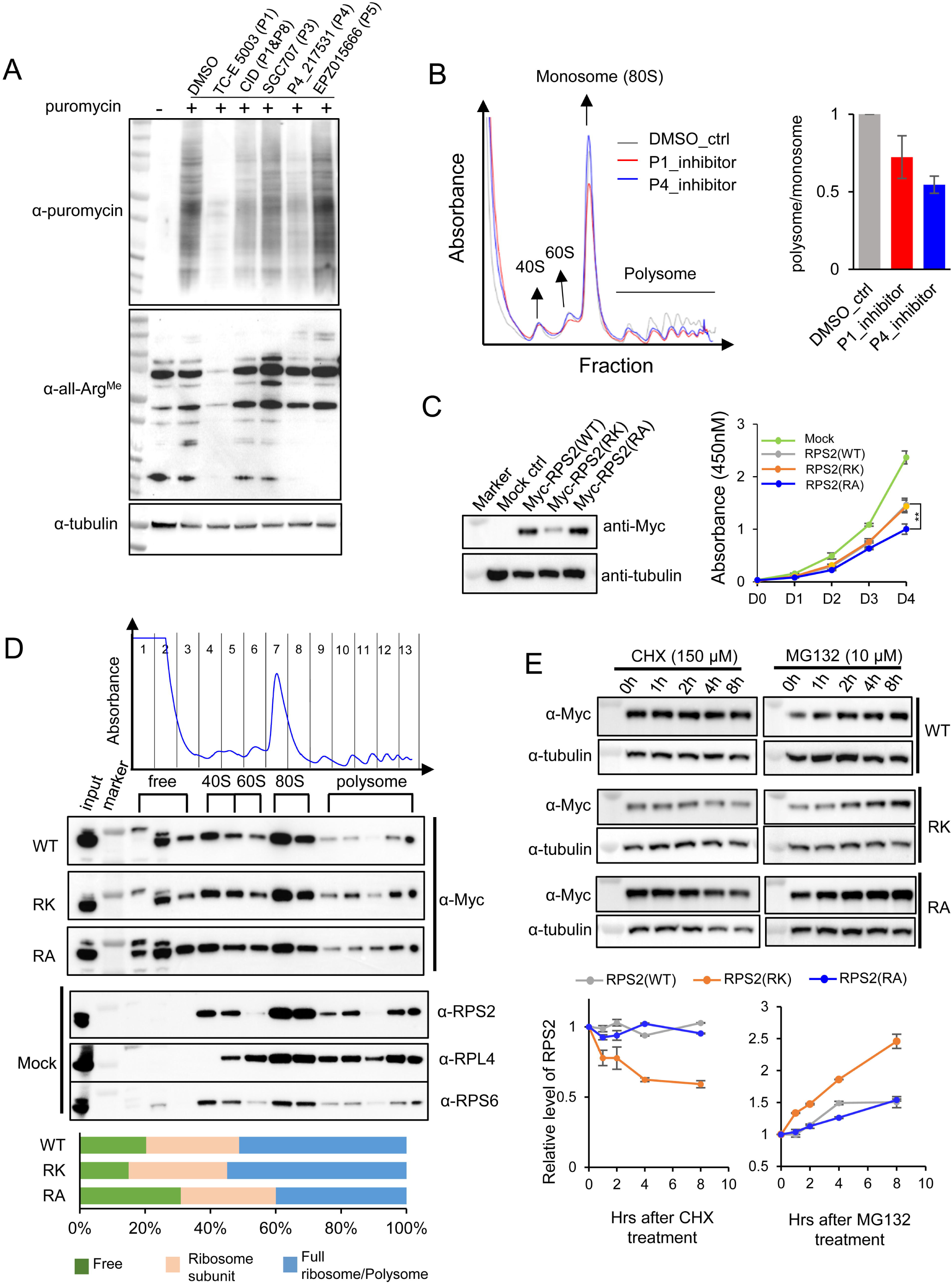
Arginine methylation affects translation efficiency and assembly of ribosomal proteins. (A) Puromycin incorporation assay of translation efficiency upon inhibition of specific PRTMs. HEK293T cells were treated with different PRMT inhibitors for 24h, and puromycin was added 30 min before cell harvest. The incorporations of puromycin were detected by western blot using anti-puromycin antibody. The pan-methylarginine antibody was used to measure changes in arginine methylation status. (B) The effect of PRMT inhibition on ribosome fractions. The HEK293T cells treated with PRMT1 and PRMT4 inhibitors were analyzed using polysome profiling, with the percent of monosome *vs.* polysomes calculated by the peak areas. In the two biological replicas, the percentage in each drug-treated sample was normalized to DMSO control. (C) Expression of Myc-tagged wild-type or mutated RPS2 in HEK293T cells, cell proliferation was followed for four days after 24hr transfection, two biological replicates were performed and ** indicates the p-value is less than 0.01. (D) Arginine methylation of RPS2 affects ribosome assembly. Myc-RPS2(WT), Myc-RPS2(6RK) and Myc-RPS2(6RA) were transfected into HEK293T cells for 48hr, followed by polysome profiling. Fractions were collected and the assembly of RPS2 were detected using western blotting with anti-Myc. The mock control fractions were also detected with anti-RPS2. Anti-RPS6 and anti-RPL4 antibodies were applied as ribosomal profiling markers. (E) Measurement of protein degradation on wild type and mutated RPS2 after transfection and treatment by cycloheximide (CHX) or MG132. The HEK293T cells were transfected with equal amount of Myc-RPS2(WT), Myc-RPS2(6RK) and Myc-RPS2(6RA), and the drugs were added 36hrs after transfection. The protein levels were detected by western blotting at different hours upon drug treatment.

We found that the inhibitors of PRMT1 and PRMT4/CARM1 effectively reduced arginine methylation and global protein synthesis across different cell lines (Fig. 6A and Fig. S7A), and the inhibition of these two PRMTs produced the most obvious reduction in arginine methylation. Using polysome profiling, we found that the inhibition of PRMT1 or PRMT4 effectively reduced the abundances of polysomes *vs.* monosomes (Fig. 6B and Fig. S7A), suggesting a global reduction of mRNAs undergoing active translation. As expected, we found that the treatment with PRMT1 or PRMT4 inhibitors dramatically slowed down cell growth, consistent with the general inhibition of protein translation (Fig. S7B-S7C).

To further examine the potential mechanisms of how arginine methylation affects mRNA translation, we selected the ribosomal protein RPS2, a newly identified PRMT1 and PRMT4 substrate in our dataset, for more detailed study. RPS2 has an N-terminal GR-rich motif that is the consensus motif for efficient arginine methylation. We made two mutations on the potential methylarginine sites (6RA and 6RK, with 6 Arg substituted by Ala or Lys respectively, see table S2) to examine if such mutations can affect the assembly of RPS2 into ribosome. We expressed the wild type and mutated RPS2 in 293 cells (Fig. 6C, left), and found that the cell proliferation were slightly delayed after ectopic expression of RPS2 (Fig. 6C, right), probably because the alteration of riboprotein ratio disrupted ribosomal assembly/function ^45^.

We further used polysome profiling to determine the efficiency of different RPS2s in assembling into ribosomal complex. While the RPS2 was mostly assembled into either ribosomal subunit or full ribosome/polysome in the control cells, a notable fraction of ectopic expressed PRS2s were found in the fraction of free proteins (Fig. 6D and Fig. S8). We found that compared to the wild type, the RPS2-RA mutation with defected methylation sites showed an assembly defect, with more proteins found in the free fraction. Interestingly, the RPS2-RK mutation showed an opposite effect, with less protein in free fraction and more in the full ribosome/polysome (Fig. 6D). Since Lysine residues can undergo various PTMs including methylation and ubiquitination, we speculate that the phenotype of RPS2-RK mutation may also be influenced by other factors. Notably, the RPS-RK mutation was expressed in a lower level compared to wild type RPS2 or RA mutation (Fig. 6C). When inhibiting translation with cycloheximide (CHX), we found that the RPS-RK is degraded more rapidly than the wild type RPS2 and RA mutation that were fairly stable (Fig. 6E). Consistently, the inhibition with MG132 significantly increased the level of RPS2-RK compared to WT and RA mutation, suggesting that RPS2-RK is degraded through ubiquitin-dependent pathway (Fig. 6E).

### Inhibition of PRMT activity cause global translation deficiency

To further examine how PRMT inhibitions affect the translation of different mRNAs, we sequenced the mRNA population associated with different ribosomal fractions after treatment of PRMT1 and PRMT4 inhibitors (Fig. 7A). Compared to the control samples treated with DSMO, the inhibition of PRMTs can significantly reduce the level of mRNAs bound by single ribosomes or polysomes (Fig. 7B). More specifically, among the 6783 protein-coding genes detected with reliable numbers of RNA-seq reads, 4392 mRNAs in the PRMT1-inhibited sample (∼65%) and 3830 mRNAs in the PRMT4-inhibited sample (∼56%) showed a consistent decrease in the association with different ribosomal fractions (i.e.,both monosome and polysomes), suggesting a general reduction of translation efficiency on most mRNAs (Fig. 7C). Interestingly, the GO analysis of these mRNAs failed to produce any functional enrichment (not shown), again suggesting a global reduction of mRNA translation rather than translation inhibition on a specific subgroup of mRNAs.

**Figure 7.**
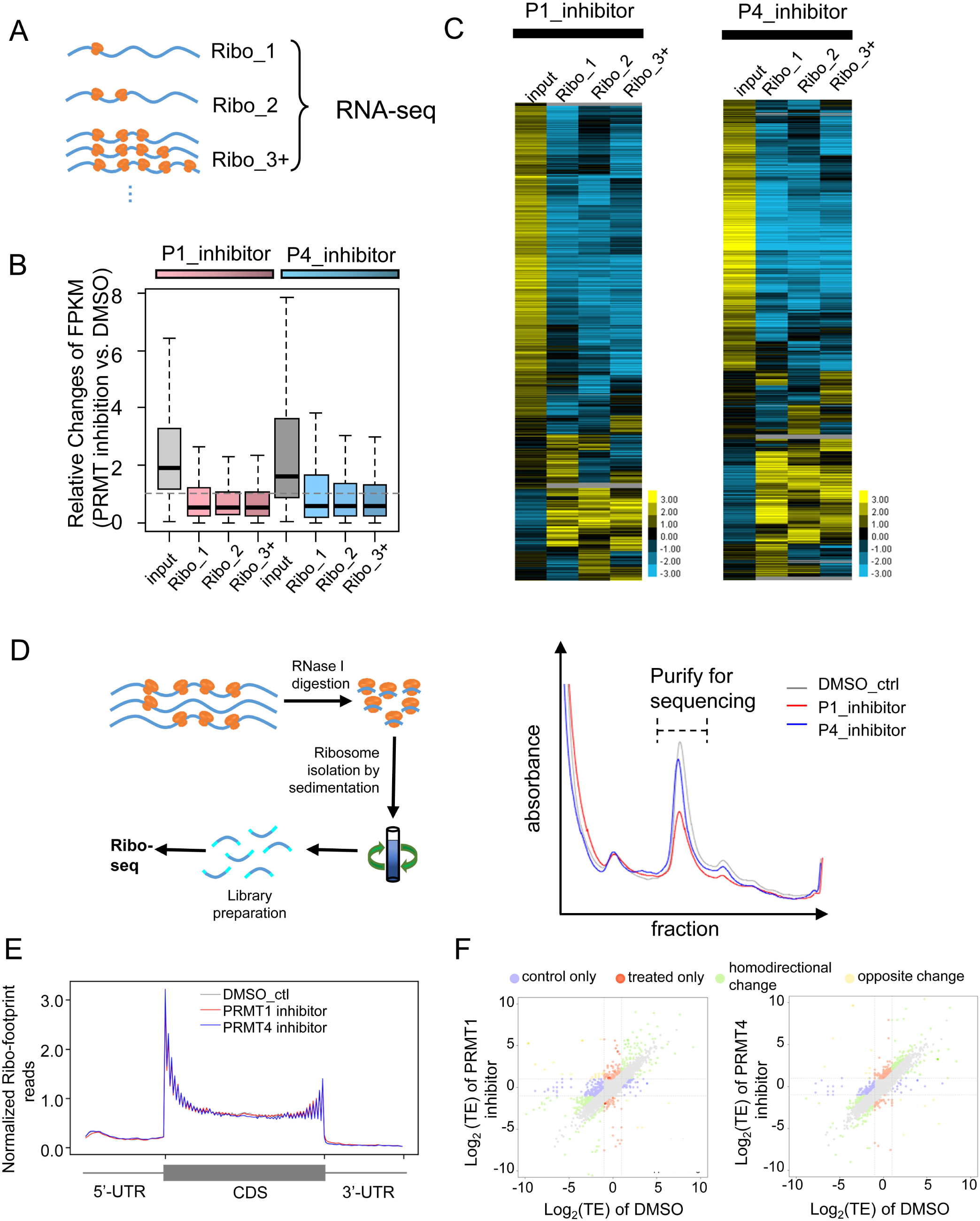
The inhibition of PRMT activity leads to global translation deficiency in thousands of genes. (A) Schematic diagram of experiments. Polysome profiling was performed to fractionate different ribosome fractions upon treatment of PRMT inhibitors in HEK293T cells. The mRNAs bound to one ribosome (ribo_1), two ribosome (ribo_2) and more than three ribosomes (ribo_3+) were collected and subjected to RNA-seq, respectively. (B) The relative FPKM changes of input mRNAs and ribosome-bound mRNAs (with PRMT1 and PRMT4 inhibition compared to DMSO control) were represented as box plot. (C) Hierarchical clustering of different mRNAs in the input and ribosome-bound fractions after treatment with PRMT1 and PRMT4 inhibitors. The log2 fold change of each mRNA was calculated and hierarchically clustered. (D) The samples with PRMT1 and PRMT4 inhibition were treated with RNase I to collect ribosome protected RNAs for high-throughput sequencing (Ribo-seq assay). The schematic diagram of experiments (left) and the isolated ribosome fractions for sequencing (right) were shown. (E) Ribosome protected RNA reads were mapped to the human genome, with the number of ribosome footprint reads in the different region of transcripts being normalized by average coverage of each transcript. All transcripts were combined to plot the distribution of normalized reads along the transcript regions. (F) Changes of translation efficiency (TE) upon PRMT inhibition. The changes of each transcript were plotted as scatter plot (more significant p-values presented with darker color). Blue: genes with large TE changes only in control sample; Red: genes with large TE changes in PRMT1 or PRMT4 inhibition sample; Green: genes with TE changes homodirectionally in two conditions; Yellow: genes with TE changes oppositely in two conditions.

We next determined how PRMT inhibition affects translation of different mRNAs using Ribo-seq to measure the distribution of ribosome protected RNA fragments in control and PRMT inhibition samples (Fig. 7D). This analysis can generate a “snapshot” of all mRNAs that are occupied by active ribosomes (i.e., undergoing active translation) in a cell at a particular condition ^46^. As expected, the ribosome occupancy is higher in the coding region compared to the 5’ and 3’ UTRs upon normalized against average coverage (Fig. 7E). In addition, the binding of ribosomes on mRNA was slightly enriched in the region around the start codon and before the stop codon, suggesting a ribosome pausing after the initiation and the delayed ribosome release (Fig. 7E), which is also consistent with the ribosomal profiling results from other groups (reviewed in 46). Interestingly, we found that the inhibition of PRMT activity did not change the distribution of ribosome occupancy on different regions of mRNAs (Fig. 7E). Given the observation of translation reduction by PRMT inhibitions (Fig. 6A and 6B), this result again suggested that PRMTs may affect the maturation of ribosomes before they are assembled onto the mRNAs, consistent with our finding that arginine methylation of ribosomal proteins is essential for ribosome assembly.

We further determined the specific genes whose translation efficiency (TE) was preferably affected by PRMT inhibition by using Xtail pipeline ^47^ to measure the genes with significant TE change after PRMT inhibition (Fig. 7F). We found that the inhibition of PRMT1 significantly changed the TE of only eight coding genes, whereas the TE of 46 protein-coding genes was significantly altered upon inhibition of PRMT4 (Fig. 7F). Interestingly, among the 46 genes affected by PRMT4 inhibition, 25 ribosomal protein genes have significantly increased TE, suggesting a potential functional complementation after the translation suppression.

## Discussion

As a common but relatively underappreciated PTM, the methylation of arginine has been found in many proteins with global identifications of methylation sites ^5, 7, 48^. Many PRMTs were found to catalyze this type of PTM, however the relationship between PRMTs and their substrates were not established on a global scale. In this study, we have for the first time identified the putative substrates for each of the human PRMTs and further characterized the novel consensus methylation motifs for individual human PRMT. We found a high degree of overlap in substrate specificity of different PRMTs, as well as a significant enrichment for RNA binding proteins in the substrates of all PRMTs. In particular, the splicing factors and ribosomal proteins are heavily methylated and overrepresented in PRMT substrates, and consistently the inhibition of PRMTs leads to global deficiency of RNA translation. Collectively, the identification and characterization of substrates for all human PRMTs provide a foundation for further studies on their biological functions.

One interesting observation is that the majority of the consensus motifs for arginine methylation are short fragments with low sequence complexity, including the well known GR-rich motifs and newly identified SR- and ER-rich motifs (Fig. 2 and Fig. S2). Since low complexity domains (e.g., GR and SR-rich domain in RNA binding proteins) usually form a non-structural region, the recognition by PRMTs likely happens in the unstructured regions of proteins. This observation therefore suggests a structure independent recognition, which is supportive to the promiscuous binding between PRMTs and many of their targets. This promiscuous binding may help to explain the high overlaps between the binding partners of different PRMTs, suggesting a certain degree of functional complementation among PRMTs. Consistently, the knockout mice of most PRMTs have only mild phenotypes ^14, 49, 50^, with the exception of PRMT1 and PRMT5, which cause lethal phenotype after knockout ^51-53^. Therefore we speculate that the additional specificity is provided by the spatial/temporal control of expression for PRMTs and their potential targets.

Arginine methylation usually increases protein hydrophobicity, thus may affect how proteins interact with their partners and assemble into a functional complex. Here we found that the core ribosomal proteins are among the largest protein groups recognized and methylated by PRMTs, raising the possibility that the methylarginine modification of ribosomal proteins can affect the assembly and function of ribosomes. It is well known that ribosome heterogeneity contributes to the regulation of mRNA translation ^54, 55^, and thus we expect that the methylarginine modification status of ribosomal proteins is a major source for ribosome heterogeneity. Here we showed that mutations on methylation sites of RPS2 can inhibit its assembly into ribosomes, and found that inhibitions of certain PRMTs impose a global suppression on translation. Rather than affecting a specific step of translation, our data implied that the translation reduction may be caused by the defects of ribosome biogenesis before they are assembled onto mRNA.

Although the ribosomal proteins are significantly enriched with arginine residue and are the most overrepresented targets of PRMTs, we speculate that they are differentially modified by different PRMTs. Consequently, inhibition of different PRMTs affects the translation efficiency of distinct sets of mRNAs. More detailed analyses on how each PRMT differentially affects the assembly and functions of ribosomes will be an important subject for future studies.

Like many PTM, methylation of arginine also has the specific “readers”, “erasers” and “readers”. Although nine PRMTs were identified as methylarginine writer, so far there is only one “eraser protein”, JMJD6, was reported for methylarginine ^56^ and several proteins containing “Tudor” domains were proposed function as putative “reader” ^57-59^. We expect that the biological functions of methylarginine modification are probably determined by the networks consisting of different “writers”, “erasers”, “readers” and their substrates. Therefore, mapping such interacting network will provide useful information on the function of arginine methylation in various proteins. This study represents a start point for a comprehensive mapping of a network containing methylarginine “writers”, “erasers”, “readers” and their substrates, and thus may serve as a foundation and reference for future research on this topic.

## Materials and Methods (see supplemental methods for more detailed information)

### Resources

#### Antibodies

Detailed information for antibodies applied in this study is listed in Table S3.

#### Cell lines

HEK293T, U2OS and PC9 cells were cultured according to instructions of American Type Culture Collection (ATCC). The cell lines have been authenticated in GENEWIZ and has been tested to have no mycoplasma contamination by mycoplasma contamination test kit (C0296, Beyotime).

#### Tools

software, databases and services were available in supplemental methods.

### Plasmids

For identification of interacting proteins of each PRMT, Human PRMT1-PRMT9 were amplified by PCR and inserted into the pcDNA3.1-BirA-HA plasmid (#36047, Addgene). For *in vitro* methylation, Myc-tagged PRMTs (PRMT1 to PRMT9) were inserted into pcDNA3.1-Myc-tag plasmids. For ribosome assembly, Myc-tagged hRPS2 were generated by PCR from human cDNA and inserted in frame with pcDNA3.1-Myc. The RPS2 mutants with arginine to alanine or lysine substitution (R22/26/34/36/227/279A, 6RA and R22/26/34/36/227/279K, 6RK,) were generated by site directed mutagenesis.

### Identification of interacting proteins for each PRMT

Human PRMT1-PRMT9 were amplified by PCR and inserted into the pcDNA3.1-BirA plasmid. The BioID experiments were performed as described in Roux et al. ^21^ with minor modifications. Briefly, the HEK293T cells transiently expressing PRMT-BirA fusion proteins were collected and lysed, and the protein complexes were purified using streptavidin beads. The resulting protein mixture was further separated through HPLC using a homemade 15 cm-long pulled-tip analytical column, and analyzed using mass spectrometry. The acquired MS/MS data were compared to the UniProt database using Integrated Proteomics Pipeline. A decoy database containing the reversed sequences of all the proteins was appended to the target database to accurately estimate peptide probabilities and false discovery rate (FDR), and FDR was set at 0.01.

Three biological replicates were performed for mock control and each PRMT individually. The label free quantification (LFQ) of the proteins was carried out using the built-in algorithm maxLFQ with the “delayed normalization” option ^60^. All proteins with at least two valid LFQ intensity values in three replicates were considered, and the average intensity were subsequently calculated. We identified putative interactors of each PRMT by evaluating protein quantification values of all proteins, and scored the positive hits using an LFQ intensity cutoff of two-fold higher than the mock control.

### Motif enrichment analysis

We retrieved the full sequences of all identified interactors of each PRMT from the UniProt database. We counted all tetrapeptide with arginine amino acid at each position (candidate PRMT’s binding sites) and calculated the frequency of each tetrapeptide in each PRMT interactome and compared with the background tetrapeptide frequency of all human proteins from UniProt database. The enrichment score of each tetrapeptide was calculated as Z score based on published methods ^61^. We collected all motifs with enrichment score larger than 4 and motif number larger than 6 as an input of clustalw2 (v2.0.9) to generate a phylogenetic tree, then clustered these motifs based on branch length and modified manually to ensure the similar motifs in one class. Finally, we used Weblogo3 (WebLogo: A sequence logo generator) to draw the consensus sequence of each cluster.

### *In vitro* methylation and MS detection of arginine methylated peptides

The HEK293T cells transiently overexpressing myc-tagged PRMT were lysed, and the overexpressed proteins were purified using Anti-myc-tag Magnetic Beads (Pierce). *In vitro* methylation assay was carried out according to Cheng et al ^62^ with minor modifications. Peptide substrates containing predicted motifs and recombinant enzyme (on beads) were incubated in the presence of S-Adenosyl-L-methionine (AdoMet). The reaction mixture was further separated through HPLC using a homemade analytical column and analyzed using mass spectrometry. The acquired MS/MS data were analyzed on a homemade database including all target peptides using PEAKS (version 8.5). Methylation and dimethylation were set as a dynamic modification with mass shift at 14.01565 and 28.0313, respectively. For each peptide, the sum of the peak areas from the TIC values of the modified peptides was divided by the peak area of the total peptides and this value was used as a relative index of the methylation percentage.

### Generation of stable cell lines

Production of lentivirus was carried out according to Addgene pLKO.1 protocol. Scramble shRNA and PRMT shRNA sequences were listed in Table S3. Lentiviruses were packaged by transfection of three plasmids (pLKO.1, psPAX2, and pMD2.G.) into HEK293T cells, and the stably transfected cells were selected with puromycin for at least two weeks. The knockdown efficiency was determined by PRMT antibodies.

### RNA-seq

HEK293T cells stably transfected with scramble shRNA or shRNAs against PRMT were harvested in Trizol reagent and RNAs were extracted according to the manufacturer’s protocol. Poly(A)+ RNA-seq libraries were prepared by using Illumina TruSeq Stranded mRNA LT Sample Prep Kit (Illumina) and subjected to deep sequencing with Illumina Hiseq X10 under PE150 sequencing model.

### Immunoprecipitation and substrate validation

The lysate of HEK293T cells was separated into trisection and incubated with ADme-, SDme- and Mme-arginine antibody, respectively. The A/G PLUS-Agarose beads (Santa Cruz) were used for immunoprecipitation of methylarginine-containing substrates. The candidate substrates (ribosomal proteins) were detected via western blot.

### Polysome profiling

Polysome profiling was carried out according to established protocols ^63, 64^. DMSO or Inhibitor-treated HEK293T cells were lysed in polysome lysis buffer. The lysates were loaded onto 10-50% sucrose gradients and ultra-centrifuged. Fractions were collected using a Brandel Density Gradient Fractionation System. Protein samples could either be precipitated with Methanol/chloroform method according to Sucrose Gradient Separation Protocol (http://www.mitosciences.com/PDF/sg.pdf) or can directly undergo western blotting. The protein precipitate was assayed by western blot to detect arginine methylation status using indicated antibodies (antibodies were listed in Table S3).

### RNA-seq of polysome profiling fractions

mRNAs from indicated fractions of polysome profiling samples were extracted with TriZol reagent. RNA-seq libraries were prepared by NEBNext®EUltra™ II Directional RNA Library Prep Kit for Illumina (NEB) and subjected to deep sequencing with Illumina Hiseq X10 under PE150 sequencing model.

### Measurement of global protein synthesis by puromycin incorporation

HEK293T, U2OS or PC9 cells were incubated with specific inhibitors (see Table S3) against several PRMTs. Subsequently, puromycin incorporation assay was performed according to Kelleher et al. ^65^. Puromycin was added to the medium of inhibitor-treated cells 30 min before harvest. An equal quantity of protein lysates was separated on SDS/PAGE and probed with anti-puromycin antibody (Millipore).

### Ribosome footprint

Cleared cell lysates from polysome profiling procedure were treated with RNase I to obtain ribosome-protected mRNA fragments (RPF). Subsequently, lysates were loaded onto 10-50% sucrose gradients, ultracentrifuged and fractionated as described above. Fractions containing monoribosome particles were combined and undergone RNA clean-up by TriZol reagent. The RNA sequencing library was prepared according to Ingolia et al. ^66^ with some modifications. The RPF library was prepared as described in Illumina Small RNA Library Prep Reference Guide. RNA samples were reverse-transcribed and cDNA libraries were gel purified and amplified by limited-cycle PCR with index primers. Libraries were cleaned up and subjected to next-generation sequencing on Illumina Hiseq X10.

### Bioinformatics analyses

The R package GeneOverlap was used to test the significance of substrates overlap between different PRMTs, with total number of interacting proteins identified in this study as the background. Gene Ontology (GO) analysis of putative PRMT substrates was performed using Database for Annotation, Visualization and Integrated Discovery (DAVID, v6.8), with total proteins in human genome as background. Protein-protein interactions were obtained from STRING database ^67^ with interaction score set to high confidence, then clustered ^68^ and demonstrated in Gephi (https://gephi.org/).

For analysis of alternative splicing, the RNA-seq reads were mapped onto the human genome reference (Ensembl GRCh37), and the PSI (Percent Spliced In) values were estimated using MISO and rMATs for each annotated splicing event. For significant change of spicing were filtered using FDR cutoff of 0.01, we also required the ΔPSI cutoff at 0.1 with minimal read count at 50.

To analyze RNA-seq data after the polysome profiling, we trimmed the adaptors and low-quality bases of paired-end 150bp reads using Cutadapt (v1.18). The trimmed reads with length < 20 nt were excluded, and the remaining reads were mapped to the human genome (GRCh37 with annotation of GENCODE v27lift37) using STAR (v2.5.3a). Genes expression levels (FPKM) were estimated by RSEM, and the relative fold changes were calculated. The Hierarchical clustering of log2 fold changes was carried out using Cluster 3.0 with centered correlation and average linkage parameter, the heatmap was visualized by TreeView.

Ribo-seq data were analyzed according to Calviello et al. ^69^. The translation efficiency of each gene was estimated by dividing the TPM of ribosome-protected mRNA with the relative transcript abundance. For coverage plot, we scaled each transcript and divided 5’-UTR, CDS, and 3’-UTR regions to 20, 100, and 50 windows, respectively. The average coverage in each window was normalized to mean coverage of the entire transcript. To assess the statistical changes of translation efficiency, Ribo-seq signals and RNA-seq signals were analyzed using Xtail pipeline ^47^, and the genes with adjusted *p*-values (less than 0.05) were used as differential translation efficiency genes.

## Acknowledgements

We thank Han Yan at Omics Core of Bio-Med Big Data Center, CAS-MPG Partner Institute for Computational Biology (PICB), for assistance with next-generation sequencing. We also thank Yu-Jie Chen at Uli Schwarz Quantitative Biology Core Facility, PICB, for experimental support with this study. We thank Prof. YanZhong Yang and Prof. Mark T. Bedford at the University of Texas MD Anderson Cancer Center for their generous gift of several PRMT plasmids.

## Declarations

### Ethics approval and consent to participate

There is no human participants or animal models used in this work. All the experiments using biological samples are conducted according to the regulation of biosafety laws in China.

### Consent for publication

Not applicable

### Availability of data and material

All sequencing data during the current study are available in The National Omics Data Encyclopedia (NODE) data depository (http://www.biosino.org/node/project/detail/OEP000307), with open access after publication.

Antibodies and other resources used in this study were listed in Table S3.

### Authors’ contributions

H.H.W. and Z.W. conceived the project. H.H.W. and M.G. carried out the molecular and biochemical experiments. X.J.F., Y.H., and Z.Y.F analyzed the RNA-seq and Ribo-seq data as well as did other bioinformatic analyses. P.W., S.X.G., X.X.T, and C.P. conducted mass spectrometry experiments. X.J.F., Y. H., Y.Y., H.H.W, and Z.W. were responsible for study design and interpretation of data. All authors were involved in drafting the manuscript and revising it critically for important intellectual content.

## References

1 Paik WK, Kim S. Omega-N-methylarginine in protein. The Journal of biological chemistry 1970; 245:88–92.

2 Baldwin GS, Carnegie PR. Specific enzymic methylation of an arginine in the experimental allergic encephalomyelitis protein from human myelin. Science 1971; 171:579–581.

3 Kakimoto Y. Methylation of arginine and lysine residues of cerebral proteins. Biochimica et biophysica acta 1971; 243:31–37.

4 Gary JD, Clarke S. RNA and protein interactions modulated by protein arginine methylation. Prog Nucleic Acid Re 1998; 61:65–131.

5 Ong SE, Mittler G, Mann M. Identifying and quantifying in vivo methylation sites by heavy methyl SILAC. Nat Methods 2004; 1:119–126.

6 Bedford MT, Richard S. Arginine methylation an emerging regulator of protein function. Molecular cell 2005; 18:263–272.

7 Pahlich S, Zakaryan RP, Gehring H. Protein arginine methylation: Cellular functions and methods of analysis. Biochimica et biophysica acta 2006; 1764:1890–1903.

8 Bedford MT. Arginine methylation at a glance. J Cell Sci 2007; 120:4243–4246.

9 Blanc RS, Richard S. Arginine Methylation: The Coming of Age. Molecular cell 2017; 65:8–24.

10 Sylvestersen KB, Horn H, Jungmichel S, Jensen LJ, Nielsen ML. Proteomic Analysis of Arginine Methylation Sites in Human Cells Reveals Dynamic Regulation During Transcriptional Arrest. Molecular & Cellular Proteomics 2014; 13:2072–2088.

11 Peng C, Wong CC. The story of protein arginine methylation: characterization, regulation, and function. Expert Rev Proteomics 2017; 14:157–170.

12 Chong PA, Vernon RM, Forman-Kay JD. RGG/RG Motif Regions in RNA Binding and Phase Separation. J Mol Biol 2018; 430:4650–4665.

13 Lorton BM, Shechter D. Cellular consequences of arginine methylation. Cellular and molecular life sciences : CMLS 2019.

14 Yang Y, Bedford MT. Protein arginine methyltransferases and cancer. Nat Rev Cancer 2013; 13:37–50.

15 Poulard C, Corbo L, Le Romancer M. Protein arginine methylation/demethylation and cancer. Oncotarget 2016.

16 Stuhlinger MC, Tsao PS, Her JH, Kimoto M, Balint RF, Cooke JP. Homocysteine impairs the nitric oxide synthase pathway: role of asymmetric dimethylarginine. Circulation 2001; 104:2569–2575.

17 Larsen SC, Sylvestersen KB, Mund A et al. Proteome-wide analysis of arginine monomethylation reveals widespread occurrence in human cells. Science signaling 2016; 9:rs9.

18 Guo A, Gu H, Zhou J et al. Immunoaffinity enrichment and mass spectrometry analysis of protein methylation. Mol Cell Proteomics 2014; 13:372–387.

19 Roux KJ, Kim DI, Burke B, May DG. BioID: A Screen for Protein-Protein Interactions. Curr Protoc Protein Sci 2018; 91:19 23 11–19 23 15.

20 Roux KJ, Kim DI, Burke B. BioID: a screen for protein-protein interactions. Curr Protoc Protein Sci 2013; 74:Unit 19 23.

21 Roux KJ, Kim DI, Raida M, Burke B. A promiscuous biotin ligase fusion protein identifies proximal and interacting proteins in mammalian cells. Journal of Cell Biology 2012; 196:801–810.

22 Shishkova E, Zeng H, Liu F et al. Global mapping of CARM1 substrates defines enzyme specificity and substrate recognition. Nature communications 2017; 8:15571.

23 Goulet I, Gauvin G, Boisvenue S, Cote J. Alternative splicing yields protein arginine methyltransferase 1 isoforms with distinct activity, substrate specificity, and subcellular localization. The Journal of biological chemistry 2007; 282:33009–33021.

24 Tang J, Gary JD, Clarke S, Herschman HR. PRMT 3, a type I protein arginine N-methyltransferase that differs from PRMT1 in its oligomerization, subcellular localization, substrate specificity, and regulation. Journal of Biological Chemistry 1998; 273:16935–16945.

25 Frankel A, Clarke S. PRMT3 is a distinct member of the protein arginine N-methyltransferase family. Conferral of substrate specificity by a zinc-finger domain. The Journal of biological chemistry 2000; 275:32974–32982.

26 Saha K, Adhikary G, Eckert RL. MEP50/PRMT5 Reduces Gene Expression by Histone Arginine Methylation and this Is Reversed by PKCdelta/p38delta Signaling. The Journal of investigative dermatology 2016; 136:214–224.

27 Guderian G, Peter C, Wiesner J et al. RioK1, a new interactor of protein arginine methyltransferase 5 (PRMT5), competes with pICln for binding and modulates PRMT5 complex composition and substrate specificity. The Journal of biological chemistry 2011; 286:1976–1986.

28 Bedford MT, Clarke SG. Protein arginine methylation in mammals: who, what, and why. Molecular cell 2009; 33:1–13.

29 Thandapani P, O’Connor TR, Bailey TL, Richard S. Defining the RGG/RG motif. Molecular cell 2013; 50:613–623.

30 Cheng D, Cote J, Shaaban S, Bedford MT. The arginine methyltransferase CARM1 regulates the coupling of transcription and mRNA processing. Molecular cell 2007; 25:71–83.

31 Feng Y, Maity R, Whitelegge JP et al. Mammalian protein arginine methyltransferase 7 (PRMT7) specifically targets RXR sites in lysine- and arginine-rich regions. The Journal of biological chemistry 2013; 288:37010–37025.

32 Huang da W, Sherman BT, Lempicki RA. Bioinformatics enrichment tools: paths toward the comprehensive functional analysis of large gene lists. Nucleic acids research 2009; 37:1–13.

33 Huang da W, Sherman BT, Lempicki RA. Systematic and integrative analysis of large gene lists using DAVID bioinformatics resources. Nat Protoc 2009; 4:44–57.

34 Matera AG, Wang Z. A day in the life of the spliceosome. Nature reviews Molecular cell biology 2014; 15:108–121.

35 Kuhn P, Chumanov R, Wang Y, Ge Y, Burgess RR, Xu W. Automethylation of CARM1 allows coupling of transcription and mRNA splicing. Nucleic acids research 2011; 39:2717–2726.

36 Bezzi M, Teo SX, Muller J et al. Regulation of constitutive and alternative splicing by PRMT5 reveals a role for Mdm4 pre-mRNA in sensing defects in the spliceosomal machinery. Genes & development 2013; 27:1903–1916.

37 Chang FN, Navickas IJ, Chang CN, Dancis BM. Methylation of Ribosomal-Proteins in Hela-Cells. Archives of Biochemistry and Biophysics 1976; 172:627–633.

38 Goldenberg CJ, Eliceiri GL. Methylation of Ribosomal-Proteins in Hela-Cells. Biochimica et biophysica acta 1977; 479:220–234.

39 Kruiswijk T, Kunst A, Planta RJ, Mager WH. Modification of yeast ribosomal proteins. Methylation. The Biochemical journal 1978; 175:221–225.

40 Polevoda B, Sherman F. Methylation of proteins involved in translation. Mol Microbiol 2007; 65:590–606.

41 Shin HS, Jang CY, Kim HD, Kim TS, Kim S, Kim J. Arginine methylation of ribosomal protein S3 affects ribosome assembly. Biochemical and biophysical research communications 2009; 385:273–278.

42 Ren J, Wang Y, Liang Y, Zhang Y, Bao S, Xu Z. Methylation of ribosomal protein S10 by protein-arginine methyltransferase 5 regulates ribosome biogenesis. The Journal of biological chemistry 2010; 285:12695–12705.

43 Simsek D, Tiu GC, Flynn RA et al. The Mammalian Ribo-interactome Reveals Ribosome Functional Diversity and Heterogeneity. Cell 2017; 169:1051–1065 e1018.

44 Choi S, Jung CR, Kim JY, Im DS. PRMT3 inhibits ubiquitination of ribosomal protein S2 and together forms an active enzyme complex. Biochimica et biophysica acta 2008; 1780:1062–1069.

45 Tye BW, Commins N, Ryazanova LV et al. Proteotoxicity from aberrant ribosome biogenesis compromises cell fitness. Elife 2019; 8.

46 Ingolia NT. Ribosome Footprint Profiling of Translation throughout the Genome. Cell 2016; 165:22–33.

47 Xiao Z, Zou Q, Liu Y, Yang X. Genome-wide assessment of differential translations with ribosome profiling data. Nature communications 2016; 7:11194.

48 Boisvert FM, Cote J, Boulanger MC, Richard S. A proteomic analysis of arginine-methylated protein complexes. Mol Cell Proteomics 2003; 2:1319–1330.

49 Jeong HJ, Lee HJ, Vuong TA et al. Prmt7 Deficiency Causes Reduced Skeletal Muscle Oxidative Metabolism and Age-Related Obesity. Diabetes 2016; 65:1868–1882.

50 Penney J, Seo J, Kritskiy O et al. Loss of Protein Arginine Methyltransferase 8 Alters Synapse Composition and Function, Resulting in Behavioral Defects. J Neurosci 2017; 37:8655–8666.

51 Nicholson TB, Chen T, Richard S. The physiological and pathophysiological role of PRMT1-mediated protein arginine methylation. Pharmacol Res 2009; 60:466–474.

52 Tee WW, Pardo M, Theunissen TW et al. Prmt5 is essential for early mouse development and acts in the cytoplasm to maintain ES cell pluripotency. Genes & development 2010; 24:2772–2777.

53 Pawlak MR, Scherer CA, Chen J, Roshon MJ, Ruley HE. Arginine N-methyltransferase 1 is required for early postimplantation mouse development, but cells deficient in the enzyme are viable. Molecular and cellular biology 2000; 20:4859–4869.

54 Emmott E, Jovanovic M, Slavov N. Ribosome Stoichiometry: From Form to Function. Trends in biochemical sciences 2019; 44:95–109.

55 Genuth NR, Barna M. The Discovery of Ribosome Heterogeneity and Its Implications for Gene Regulation and Organismal Life. Molecular cell 2018; 71:364–374.

56 Chang B, Chen Y, Zhao Y, Bruick RK. JMJD6 is a histone arginine demethylase. Science 2007; 318:444–447.

57 Vagin VV, Wohlschlegel J, Qu J et al. Proteomic analysis of murine Piwi proteins reveals a role for arginine methylation in specifying interaction with Tudor family members. Genes & development 2009; 23:1749–1762.

58 Kirino Y, Vourekas A, Sayed N et al. Arginine methylation of Aubergine mediates Tudor binding and germ plasm localization. RNA 2010; 16:70–78.

59 Chen C, Nott TJ, Jin J, Pawson T. Deciphering arginine methylation: Tudor tells the tale. Nature reviews Molecular cell biology 2011; 12:629–642.

60 Cox J, Hein MY, Luber CA, Paron I, Nagaraj N, Mann M. Accurate proteome-wide label-free quantification by delayed normalization and maximal peptide ratio extraction, termed MaxLFQ. Mol Cell Proteomics 2014; 13:2513–2526.

61 Fairbrother WG, Yeh RF, Sharp PA, Burge CB. Predictive identification of exonic splicing enhancers in human genes. Science 2002; 297:1007–1013.

62 Cheng D, Vemulapalli V, Bedford MT. Methods applied to the study of protein arginine methylation. Methods Enzymol 2012; 512:71–92.

63 Lin CJ, Robert F, Sukarieh R, Michnick S, Pelletier J. The antidepressant sertraline inhibits translation initiation by curtailing mammalian target of rapamycin signaling. Cancer research 2010; 70:3199–3208.

64 Vyas K, Chaudhuri S, Leaman DW et al. Genome-wide polysome profiling reveals an inflammation-responsive posttranscriptional operon in gamma interferon-activated monocytes. Molecular and cellular biology 2009; 29:458–470.

65 Kelleher AR, Kimball SR, Dennis MD, Schilder RJ, Jefferson LS. The mTORC1 signaling repressors REDD1/2 are rapidly induced and activation of p70S6K1 by leucine is defective in skeletal muscle of an immobilized rat hindlimb. Am J Physiol-Endoc M 2013; 304:E229–E236.

66 Ingolia NT, Brar GA, Rouskin S, McGeachy AM, Weissman JS. The ribosome profiling strategy for monitoring translation in vivo by deep sequencing of ribosome-protected mRNA fragments. Nat Protoc 2012; 7:1534–1550.

67 Szklarczyk D, Franceschini A, Wyder S et al. STRING v10: protein-protein interaction networks, integrated over the tree of life. Nucleic acids research 2015; 43:D447–452.

68 Hu Y. Efficient, high-quality force-directed graph drawing. Mathematica journal 2005; 10:37–71.

69 Calviello L, Mukherjee N, Wyler E et al. Detecting actively translated open reading frames in ribosome profiling data. Nat Methods 2016; 13:165–170.

