## Supplemental methods and figures for "A systematic survey of PRMT interactomes reveals key roles of arginine methylation in the global control of RNA splicing and translation"

### Supplemental Files

- **Supplementary experimental procedures**

- **Supplemental figures**:

**Figure S1 (related to Figure 1).** Subcellular localization of PRMT-BirA* fusion protein.

**Figure S2 (related to Figure 1).** Substrate characteristics of PRMT interactome.

**Figure S3 (related to Figure 2).** The sequence clustering and consensus motifs for the putative substrate of each PRMT.

**Figure S4 (related to Figure 2).** In vitro methylation assay.

**Figure S5 (related to Figure 3).** The analyses of enriched domains in putative PRMT substrates.

**Figure S6 (related to Figure 4).** The validation of splicing changes in specific AS events.

**Figure S7 (related to Figure 6).** PRMT inhibition can lead to translation deficiency in other cell types and inhibition of cell proliferation.

**Figure S8 (related to Figure 6).** Arginine methylation of RPS2 affects ribosome assembly.

- **Supplemental tables**: Table S1-S3

**Table S1**. Interactome of each human PRMT.

**Table S2.** Core ribosomal proteins with interacting PRMTs and putative methyl-arginine sites.

**Table S3**. Antibodies and other resources used in this study.

### Supplementary experimental procedures

#### Identification of interacting proteins for each PRMT

Human PRMT1-PRMT9 were amplified by PCR and inserted in frame into the pcDNA3.1-BirA plasmid (#36047, Addgene). The BioID experiment was performed as described in Roux et al. [1] with minor modifications. In brief, pcDNA3.1-BirA plasmid (negative control) or pcDNA3.1-PRMT-BirA plasmids were transiently transfected into $5\times10^6$ 293T cells in 15 cm dishes with lipo3000 reagent (Life Technologies). Then cells were incubated in complete media supplemented with 50 μM biotin (B4639, Sigma-Aldrich) for 36hr. Subsequently, cells were collected and lysed in 1ml lysis buffer (50mM Tris-HCl, pH = 7.4, 150 mM NaCl, 1% Triton X-100, 1% deoxycholate, and 0.1% SDS (Beyotime, China) with 1×Complete protease inhibitor cocktail (Roche)). After sonication and centrifugation, supernatants were incubated with 400 μl Dynabeads (M-280 Streptavidin, Invitrogen) overnight.

The beads were collected and washed in very stringent wash buffer (2% SDS) then kept in 50 mM $NH_4HCO_3$ for mass spectrometry analysis. For the assay of input protein, 5% of the beads were taken out and assessed labeling consequences via western blot by using Streptavidin-HRP (CST) as the antibody. Another 5% of beads were assessed quality by SDS-PAGE and silver staining. For the purification of PRMT interacting proteins, the beads were mixed with 50 μl denature buffer (8 M urea in 100mM Tris-HCl, pH 8.5) and sonicated for 30 minutes. Dithiothreitol (Sigma Aldrich, final concentration is 10 mM) was added to the solution and incubated at 56 °C for 30 minutes for reduction. Subsequently, Iodoacetamide (Sigma Aldrich) was added at a final concentration of 15mM and incubated at room temperature for 30 minutes for alkylation. The protein mixture was subsequently diluted and digested with Trypsin (Promega) at 1:50 (w/w). The digestion was stopped by Formic Acid (Sigma Aldrich), and the peptide mixture was desalted by monospin C18 column (SHIMADZU-GL).

The peptide mixture was loaded onto a homemade 15 cm-long pulled-tip analytical column (aqua, C18, 750 μm OD× 360 μm ID, 3μm particle size, 125Å pore diameter, Phenomenex, Torrance, CA) connected to a nanoACQUITY UPLC (Waters) for mass spectrometry analysis. The elution gradient and mobile phase constitution used for peptide separation were as follows: 0-3 min, 3%-6% B; 3-75 min, 6-35% B; 75-100 min, 35-45% B; 100-103 min, 45-100% B; 103-113 min, 100%-100% B, 113-114 min, 100-3% B; 114-120 min, 3% B (buffer A: 0.1% Formic Acid in Water and buffer B: 0.1% formic acid in Acetonitrile) at a flow rate of 300 nl/min. Peptides eluted from the LC column were directly electrosprayed into the mass spectrometer with a distal 1.8-kV spray voltage. Survey full-scan MS spectra (from m/z 350–1800) were acquired in the Orbitrap analyzer (Q Exactive) with resolution r =70,000 at m/z 400. The top 20 MS/MS events were sequentially generated from the full MS spectrum at a 30% normalized collision energy. The acquired MS/MS data were compared to the UniProt database by Andromeda algorithm built-in MaxQuant engine (v1.5.5.0). using Integrated Proteomics Pipeline (IP2, http://integratedproteomics.com/). Mass tolerances were set at 20 ppm and 0.02 Da for precursor ions and MS/MS search, respectively. Trypsin was defined as cleavage enzyme; Cysteine alkylation by iodoacetamide was specified as fixed modification with mass shift 57.02146; Methionine oxidation and protein N-terminal biotinylation were set as dynamic modification with mass shift 15.9949 and 226.0776. A decoy database containing the reversed sequences of all the proteins was appended to the target database to accurately estimate peptide probabilities and false discovery rate (FDR), and FDR was set at 0.01. A built-in label free quantification algorithm called maxLFQ was used for peptide quantification with a delayed normalization method [2] to normalize peptide intensity quantification across all the samples.

For data reliability, we performed three biological replicates. All proteins with at least two valid LFQ intensity values in three replicates were considered, and the average intensity were subsequently calculated. We identified putative interactors of each PRMT by evaluating protein quantification values of all proteins, and scored the positive hits using an LFQ intensity cutoff of two-fold higher than the mock control. The interactome of each PRMT were listed in Table S1.

**Bioinformatics analyses**

The R package GeneOverlap was used to test the significance of substrates overlap between different PRMTs, with total number of interacting proteins identified in this study as the background. The putative PRMT substrates was performed using Database for Annotation, Visualization and Integrated Discovery (DAVID, v6.8), with total proteins in human genome as background. Protein-protein interactions were obtained from STRING database [3] (v10.5, minimum required interaction score was set to high confidence as 0.7),. then clustered [4] and demonstrated in Gephi (https://gephi.org/).

**Motif enrichment analysis：**

We retrieved the full sequences of all identified interactors of each PRMT from the UniProt database. We counted all tetrapeptide with arginine amino acid at each position (candidate PRMT's binding sites) and calculated the frequency of each tetrapeptide in each PRMT interactome and compared with the background tetrapeptide frequency of all human proteins from UniProt database. The enrichment score of each tetrapeptide was calculated as Z score based on published methods [5]. We collected all motifs with enrichment score larger than 4 and motif number larger than 6 as an input of Clustalw2 (v2.0.9) to generate a phylogenetic tree, then clustered these motifs based on branch length and modified manually to ensure the similar motifs in one class. Finally, we used Weblogo3 (WebLogo: A sequence logo generator) to draw the consensus sequence of each cluster.

***In vitro* methylation and MS detection of arginine methylated peptides**

Each Myc-tagged PRMT construct was transiently transfected into HEK293T cells in a 15 cm dish using lipo3000 reagent (Invitrogen). After 48hr expression, cell pellets were collected and lysed in RIPA buffer (20 mM Tris-HCl, pH 7.4, 150 mM NaCl, 5 mM EDTA pH 80, 1% NP-40, 0.5% deoxycholate, 0.1% SDS, supplemented with 1× protease inhibitor cocktail) with 30 min rotation at 4 ˚C. The crude lysate was cleared by centrifugation then mixed with 50 µl pre-washed anti-myc-tag Magnetic Beads (Pierce), followed by incubating at room temperature for 2h. The beads were washed four times with lysis buffer and kept in methylation buffer for further *in vitro* methylation experiment. 10% of input were taken out and assessed the purity by SDS-PAGE and Coomassie Blue staining.

The candidate peptide substrates containing predicted motifs were synthesized by GenScript. *In vitro* methylation assay was carried out according to Cheng et al [6] with minor modifications. Briefly, 0.2-0.5 µg recombinant enzyme (on beads) and 1µg peptide substrates were incubated in methylation buffer (50mM Tris-HCl, pH 8.0, 100 mM NaCl and 0.5mM EDTA) in the presence of 0.1µg S-Adenosyl-L-methionine (AdoMet, Sigma) at 30 ˚C for two hours. The reaction was stopped by adding 5% formic acid then underwent HPLC/MS as soon as possible. The reaction mixture was desalted by monospin C18 column (SHIMADZU-GL), and then loaded onto a homemade 15 cm pulled-tip analytical column (Aqua, C18, 750 µm OD × 360 µm ID, 3µm particle size, 125Å pore diameter, Phenomenex, Torrance, CA) connected to an Easy-nLC 1000 nano HPLC (Thermo Scientific, San Jose, CA) for mass spectrometry analysis. The mobile phase and elution gradient used for peptide separation were as follows: 0-1 min, 1% - 5% B; 1-15 min, 5-40% B; 15-20 min, 40% B; (mobile phase A: 0.1% FA in water and mobile phase B: 0.1% FA in Acetonitrile) at a flow rate of 300 nl/min. Peptides eluted from the LC column were directly electrosprayed into the mass spectrometer with the application of a distal 1.8-kV spray voltage. Survey full-scan MS spectra (from m/z 300–1800) were acquired in the Orbitrap analyzer with resolution r =70,000 at m/z 400. And top 20 MS/MS events were sequentially generated selected from the full MS spectrum at a 30% normalized collision energy.

The acquired MS/MS data were analyzed against a homemade database (including all target peptides) using PEAKS (version 8.5). Mass tolerances for precursor ions were set at 20 p.p.m. and were set at 0.02 Da for MS/MS. No specificity was defined as cleavage enzyme; Methylation and dimethylation were set as a dynamic modification with mass shift 14.01565 and 28.0313, respectively. For each peptide, the sum of the peak areas from the TIC values of the modified peptides was divided by the peak area of the reference unmodified peptide and this value was used as a relative index of the methylation and dimethylation.

**Generation of stable cell lines**

Production of lentivirus was carried out according to Addgene pLKO.1 protocol. Scramble shRNA and validated PRMT shRNA sequences were obtained from Sigma Aldrich and were listed in Table S3. Lentiviruses were packaged by transfection of three plasmids (pLKO.1, psPAX2, and pMD2.G.) into HEK293T cells. Supernatant was collected after 48h infection and added into HEK293T cells for transfection. The stably transfected cells were selected with 10 μg/ml puromycin for at least two weeks to obtain the stable cell lines. The knockdown efficiency was determined by PRMT antibodies (Table S3).

**RNA isolation, polyadenylated RNA separation and RNA-seq of PRMT knockdown samples**

HEK293T cells stably transfected with scramble shRNA or shRNAs against PRMT were harvested in Trizol reagent (Ambion) and RNAs were extracted according to the manufacturer's protocol. Poly(A)+ RNA-seq libraries were prepared by using Illumina TruSeq Stranded mRNA LT Sample Prep Kit (Illumina) and subjected to deep sequencing with Illumina Hiseq X10 under PE150 sequencing model.

For analysis of alternative splicing, The sequencing adapters were trimmed by Cutadapt (v1.18), then the RNA-seq reads were mapped onto the human genome reference (Ensembl GRCh37) using HISAT [7] with default parameters. The PSI (Percent-Spliced-In) values were estimated by MISO and rMATs for each annotated splicing event. For significant change of spicing were filtered using FDR cutoff of 0.01, we also required the ΔPSI cutoff at 0.1 with minimal read count at 50.

**Polysome profiling**

Polysome profiling was carried out according to Lin et al. and Vyas et al. [8,9]. Briefly, cells were grown to 70-80% confluence. After treated with cycloheximide (CHX, 100μg/ml) for 20min, cells were washed with CHX/PBS and lysed in polysome lysis buffer (100mM KCl, 5mM $MgCl_2$, 10mM HEPES, 1mM DTT, 100μg/ml CHX, supplemented with protease inhibitor, RNase inhibitor, and DNase I, pH=7.4), followed by shearing 20 times with 26-gauge needles gently. Lysates were cleared by centrifugation. 10% of cleared lysate was retained as input for total RNA and protein extraction. Half of the remained lysates were subjected to ribosome footprinting. Another half were loaded onto 12ml 10-50% sucrose gradients and ultracentrifuged for 3hr at 35,000rpm, 4˚C on an SW41Ti rotor (Beckman Coulter). 0.75ml fractions were collected using a Brandel Density Gradient Fractionation System with a 254nm filter lens. Samples from polysome profiling assay were either precipitated with Methanol/chloroform method according to Sucrose Gradient Separation Protocol (http://www.mitosciences.com/PDF/sg.pdf) or **can directly undergo western blotting without precipitation**. The samples were heated at 95 ˚C for 10 min in SDS protein sample loading buffer and assayed by western blot to detect arginine methylation status using indicated antibodies (antibodies listed in Table S3).

**Immunoprecipitation and substrate validation**

The HEK293T cell lysates were prepared using RIPA buffer (20 mM Tris-HCl, pH 7.4, 150 mM NaCl, 5 mM EDTA pH 80, 1% NP-40, 0.5% deoxycholate, 0.1% SDS, supplemented with 1× protease inhibitor cocktail). The lysate was separated into trisection and incubated with ADme-, SDme- and Mme arginine antibody, respectively, for 12 h at 4˚C. 100 μl Protein A/G PLUS-Agarose beads (Santa Cruz) were added to each sample then each mixture was incubated at 4˚C for 4 h. The beads were washed in PBS four times, followed by detection of candidate substrates (ribosomal proteins) via western blot.

**RNA-seq of polysome profiling fractions**

mRNAs from indicated fractions of polysome profiling sample were extracted with TriZol reagent. RNA-seq libraries were prepared by NEBNext® Ultra™ II Directional RNA Library Prep Kit for Illumina (NEB) after rRNA depletion with NEB E6310. The library was subsequentially subjected to deep sequencing with Illumina Hiseq X10 under PE150 sequencing model.

To analyze RNA-seq data after the polysome profiling, we trimmed the adaptors and low-quality bases of paired-end 150bp reads using Cutadapt (v1.18). The trimmed reads with length < 20 nt were excluded, and the remaining reads were mapped to the human genome (GRCh37 with annotation of gencode v27lift37) using STAR (v2.5.3a). Genes expression levels (FPKM) were estimated by RSEM, and the relative fold changes were calculated. The relative FPKM changes of input mRNAs and ribosome-bound mRNAs (with PRMT1 and PRMT4 inhibition compared to DMSO control) were represented as box plot. The Hierarchical clustering of log2 fold changes was carried out using Cluster 3.0 with centered correlation and average linkage parameter, the heatmap was visualized by TreeView.

**The measurement of global protein synthesis by puromycin incorporation**

HEK293T cells were incubated with specific inhibitors against each PRMT for 24 h (TC-E 5003 for PRMT1 (Santa Cruz), CID 5380390 for PRMT1 and PRMT8 (Sigma), 217531 for PRMT4 (Merck Millipore), EPZ015666 for PRMT5 (Sigma) and a GSK candidate inhibitor for type I PRMT (GSK023)). Subsequently, puromycin incorporation assay was performed according to Kelleher et al. [10]. Puromycin (1μM) was added to the medium of inhibitor-treated cells 30 min before harvest. Cells were lysed using RIPA buffer (see before). An equal quantity of protein lysates was separated on SDS/PAGE and probed with anti-puromycin antibody (Millipore).

**Cell growth assay by CCK8**

Mock, Myc-RPS2(WT), Myc-RPS2(6RK) and Myc-RPS2(6RA) plasmids were transfected into HEK293T cells for 24hrs. 2000 cells of each sample were counted using CountStar and seeded into 96 well-plate, was followed for four days

**Ribosome footprint and Ribo-seq analysis**

Cleared cell lysates from polysome profiling procedure were treated with RNase I (Thermo Fisher Scientific) at room temperature for 45 min to obtain ribosome-protected mRNA fragments (RPF). The reaction was stopped by adding SUPERaseIn RNase Inhibitor (Invitrogen). Subsequently, lysates were loaded onto 12ml 10-50% sucrose gradients, centrifuged and fractionated as described above. Fractions containing 80S ribosome particles were combined and underwent RNA clean-up by TriZol reagent. The RNA sequencing library was prepared according to Ingolia et al. [11] with some modifications. In brief, rRNAs were depleted in both total RNA and RPF RNA samples with Ribo-Zero Gold kit (Illumina). Total RNA was fragmented as input control. Subsequently, the ends of both the RPF RNA and fragmented total RNA samples were repaired prior to 3' and 5' adapter ligation, as described in Illumina Small RNA Library Prep Reference Guide. RNA samples were reverse-transcribed by Superscript III reverse transcriptase (Invitrogen). cDNA libraries then were gel purified and amplified by limited-cycle PCR with index primers. Libraries were cleaned up and subjected to next-generation sequencing on Illumina Hiseq X10 under PE150 sequencing model.

Ribo-seq data were analyzed according to Calviello et al. [12]. After removing adapters by Cutadapt (v1.18), reads aligning to rRNA sequences were removed using Bowtie (v1.1.2). Unaligned Ribo-seq reads and RNA-seq reads were aligned to Gencode annotation release 27lift37 (GRCh37.p13) for human using STAR (v2.4.2a) allowing a maximum of 2 mismatches and multimapping of to up to eight different positions. The translation efficiency of each gene was estimated by dividing the TPM of ribosome-protected mRNA with the relative transcript abundance from RSEM (v1.2.22). For coverage plot, we used best aligned reads to estimate the coverage of protein coding genes at each position and scaled each transcript to 20, 100, and 50 windows on 5' UTR, CDS, and 3' UTR regions, respectively. The Ribo-footprint coverage in each window was normalized to mean coverage of the entire transcript. To assess the statistical changes of translation efficiency, Ribo-seq signals and RNA-seq signals were analyzed using Xtail pipeline [13]. The genes with adjusted *p*-values (less than 0.05) were used as differential translation efficiency genes.

#### Supplementary references


1 Roux KJ, Kim DI, Raida M, Burke B. A promiscuous biotin ligase fusion protein identifies proximal and interacting proteins in mammalian cells. *Journal of Cell Biology* 2012; **196**:801-810.

2 Cox J, Hein MY, Luber CA, Paron I, Nagaraj N, Mann M. Accurate proteome-wide label-free quantification by delayed normalization and maximal peptide ratio extraction, termed MaxLFQ. *Mol Cell Proteomics* 2014; **13**:2513-2526.

3 Szklarczyk D, Franceschini A, Wyder S *et al.* STRING v10: protein-protein interaction networks, integrated over the tree of life. *Nucleic acids research* 2015; **43**:D447-452.

4 Hu Y. Efficient, high-quality force-directed graph drawing. *Mathematica journal* 2005; **10**:37-71.

5 Fairbrother WG, Yeh RF, Sharp PA, Burge CB. Predictive identification of exonic splicing enhancers in human genes. *Science* 2002; **297**:1007-1013.

6 Cheng D, Vemulapalli V, Bedford MT. Methods applied to the study of protein arginine methylation. *Methods Enzymol* 2012; **512**:71-92.

7 Kim D, Langmead B, Salzberg SL. HISAT: a fast spliced aligner with low memory requirements. *Nat Methods* 2015; **12**:357-360.

8 Lin CJ, Robert F, Sukarieh R, Michnick S, Pelletier J. The antidepressant sertraline inhibits translation initiation by curtailing mammalian target of rapamycin signaling. *Cancer research* 2010; **70**:3199-3208.
9 Vyas K, Chaudhuri S, Leaman DW *et al.* Genome-wide polysome profiling reveals an inflammation-responsive posttranscriptional operon in gamma interferon-activated monocytes. *Molecular and cellular biology* 2009; **29**:458-470.
10 Kelleher AR, Kimball SR, Dennis MD, Schilder RJ, Jefferson LS. The mTORC1 signaling repressors REDD1/2 are rapidly induced and activation of p70S6K1 by leucine is defective in skeletal muscle of an immobilized rat hindlimb. *Am J Physiol-Endoc M* 2013; **304**:E229-E236.
11 Ingolia NT, Brar GA, Rouskin S, McGeachy AM, Weissman JS. The ribosome profiling strategy for monitoring translation in vivo by deep sequencing of ribosome-protected mRNA fragments. *Nat Protoc* 2012; **7**:1534-1550.
12 Calviello L, Mukherjee N, Wyler E *et al.* Detecting actively translated open reading frames in ribosome profiling data. *Nat Methods* 2016; **13**:165-170.
13 Xiao Z, Zou Q, Liu Y, Yang X. Genome-wide assessment of differential translations with ribosome profiling data. *Nature communications* 2016; **7**:11194.

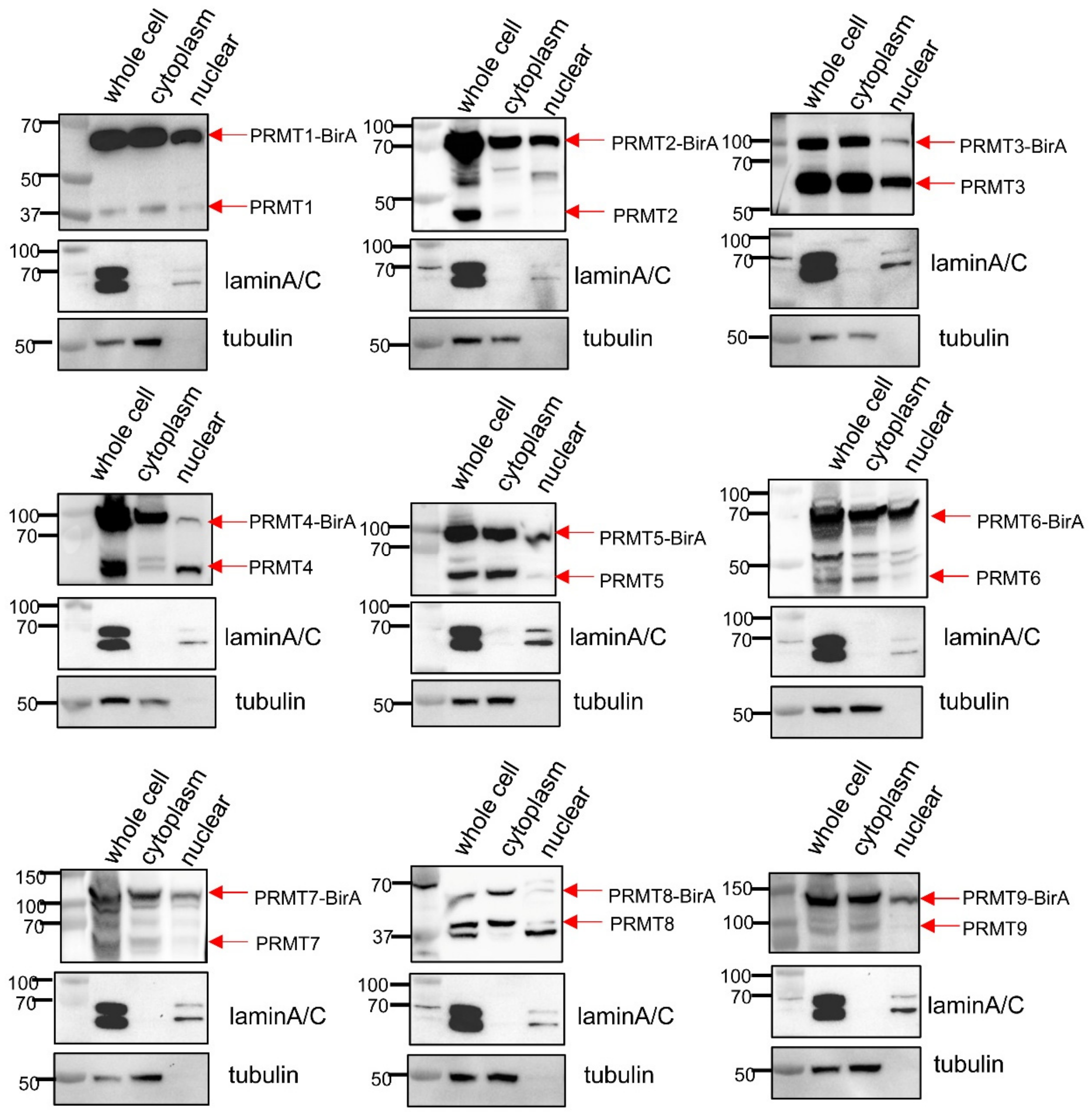


**Figure S1 (related to Figure 1). Subcellular localization of PRMT-BirA* fusion protein.** Whole lysate, cytosolic, and nuclear fractions of each PRMT-BirA* transfected HEK293T cells were prepared according to CelLytic™ NuCLEAR™ Extraction Kit (Sigma). Fractions were analyzed by Western blotting using each PRMT antibody, respectively (supplementary table 3). Lamin A/C was applied as nuclear marker and tubulin was applied as cytoplasm marker.

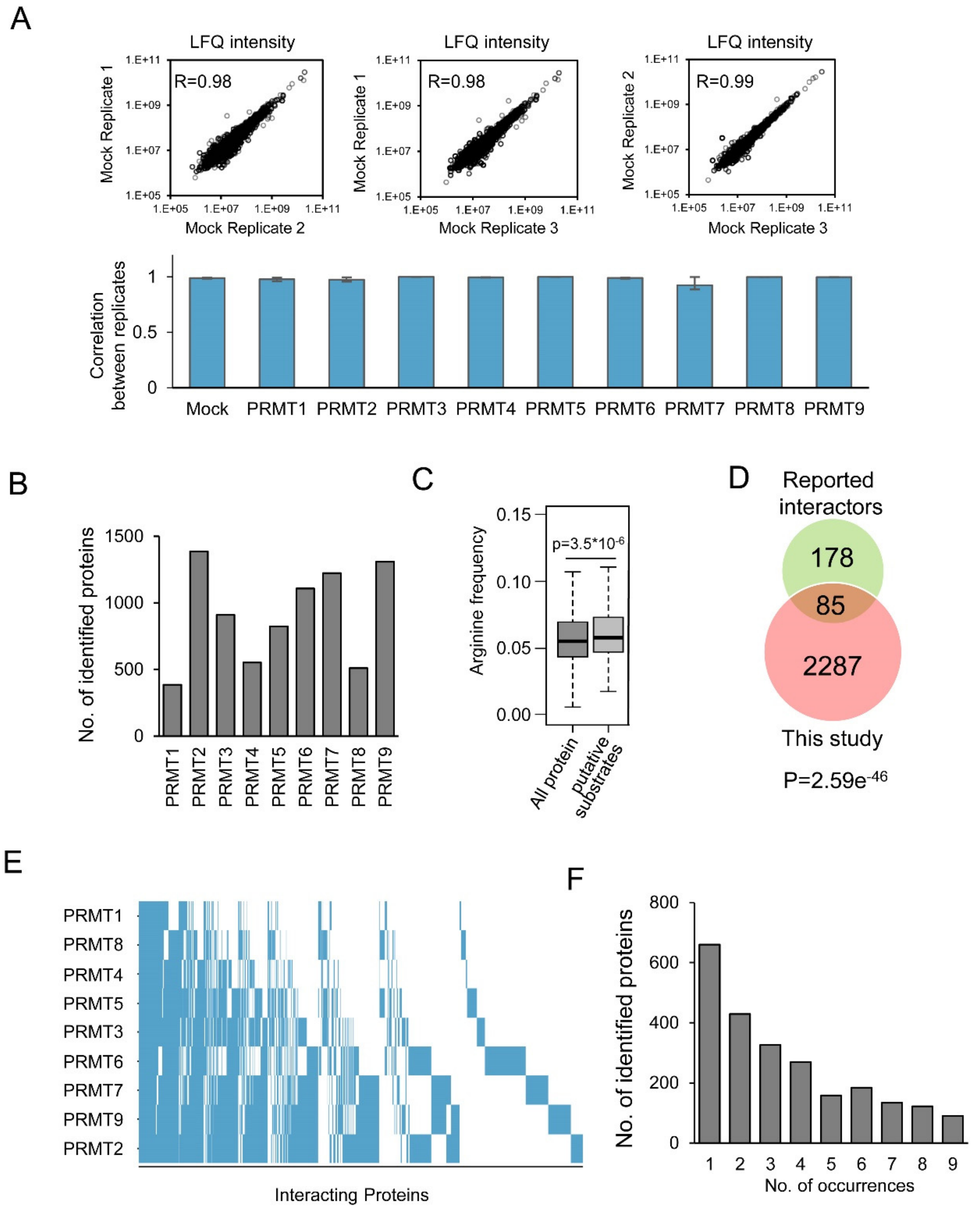


**Figure S2 (related to Figure 1). Substrate characteristics of PRMT interactome.**

(A) The LFQ intensity correlation between different replicates of mock control (upper) and all PRMTs (lower). (B) Numbers of identified interactors of each PRMT. (C) Arginine frequency of all annotated human proteins (n= 42119) and identified putative PRMT substrates (n=1416). (D) Comparison of the PRMT interactome to reported interactors listed in IntAct online PPI database (https://www.ebi.ac.uk/intact/). (E) The interacting proteins of each PRMT are listed by their shared order among nine PRMTs. (F) The number of proteins interacting with a single PRMT to multiple PRMTs in this study.

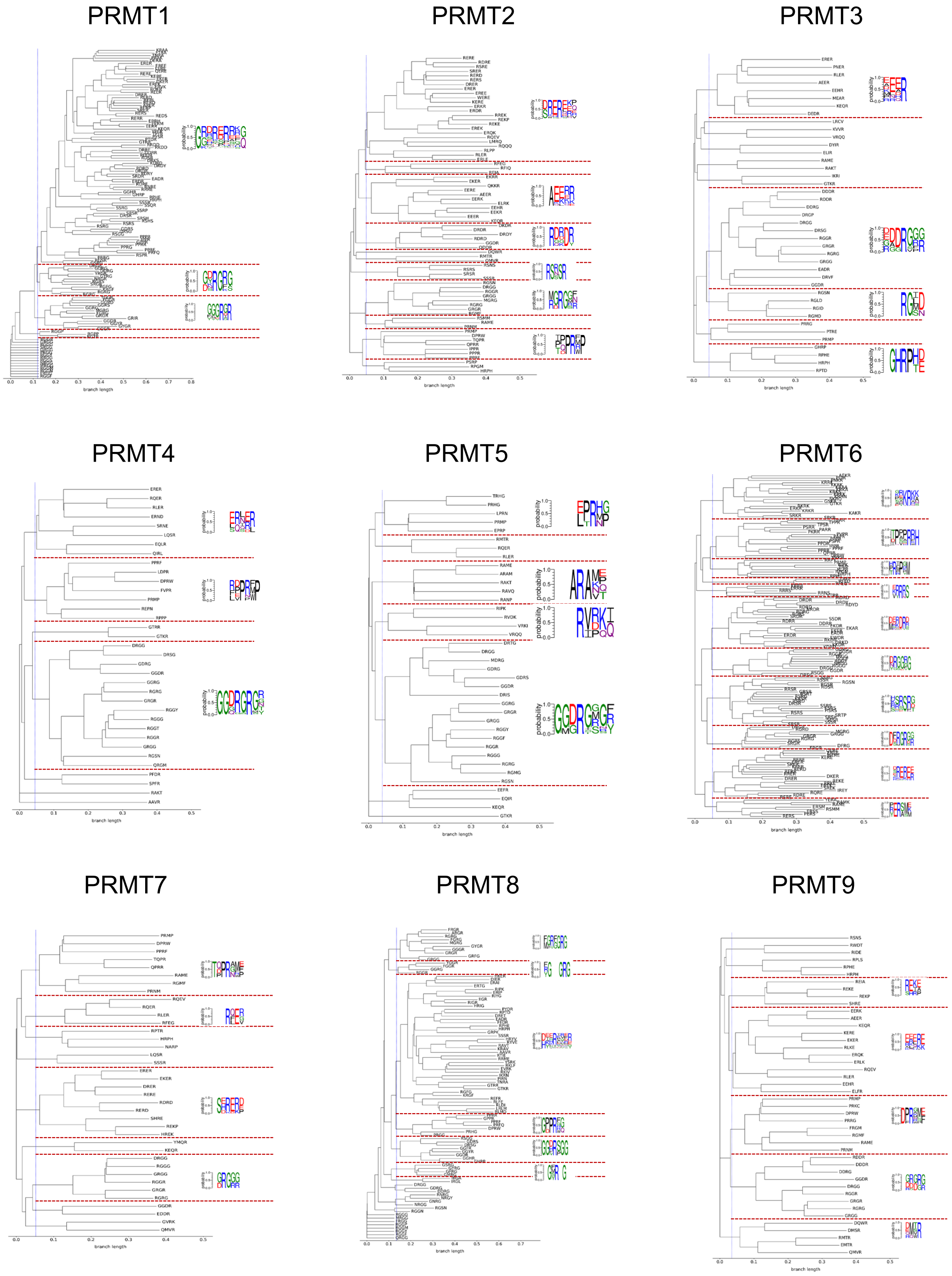


Figure S3 (related to Figure 2). The sequence clustering and consensus motifs for the putative substrate of each PRMT.

The enriched tetrapeptides in putative PRMT substrates are collected as the input of clustalw2 pipeline (v2.0.9) to generate the phylogenetic tree, and the Weblogo3 was used to generate the consensus motifs of each cluster.

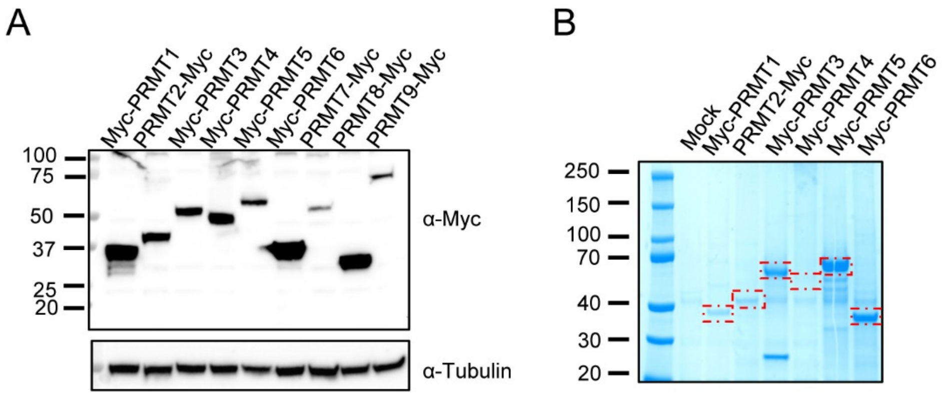


**Figure S4 (related to Figure 2). In vitro methylation assay.**

(A) Myc-tagged PRMTs were expressed in HEK293T cells and protein expression was detected by western blotting with anti-Myc-tag antibody. (B) Isolation and purification of myc-tagged PRMTs. Each Myc-tagged PRMT utilized in *in-vitro* methylation assay was purified with anti-myc magnetic beads and qualified by SDS-PAGE and Coomassie Blue staining.

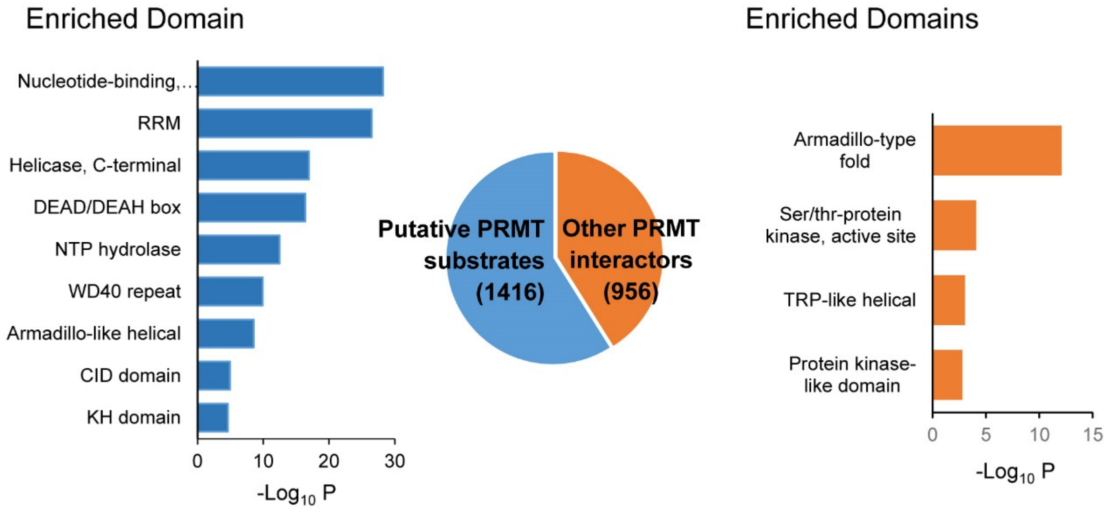


**Figure S5 (related to Figure 3). The analyses of enriched domains in putative PRMT substrates.**

The domain analysis (in DAVID) of the 1416 putative PRMT substrates detected in both this study and previously identified methylarginine-containing proteins (putative PRMT substrates, blue bars), as well as other PRMT interacting proteins identified in only this study (orange bar).

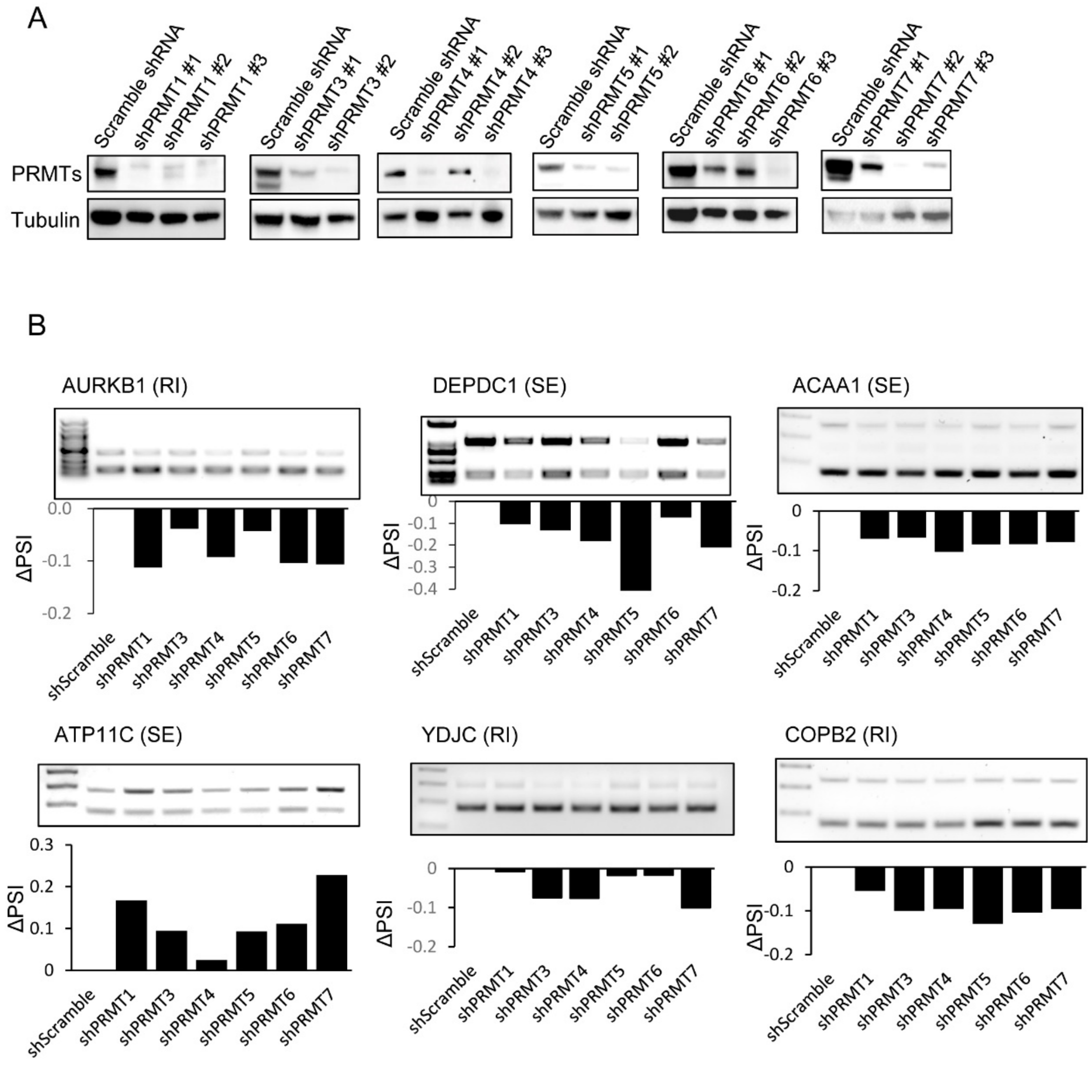


**Figure S6 (related to Figure 4). The validation of splicing changes in specific AS events.**

(A) The validation of stable knockdown against each PRMT. For each PRMT, 2 to 3 shRNA plasmids for the knockdown of each PRMT were designed and transfected into HEK293T cells to generate stable knockdown cell lines. The knockdown efficiency was validated by western blot with PRMT antibody (the knockdown against PRMT2 and PRMT9 did not work reliably using shRNA, PRMT8 does not express in HEK293T cells). (B) Validation of splicing changes in specific AS events that are altered in PRMT knockdown cells using semi-quantitative RT-PCR.

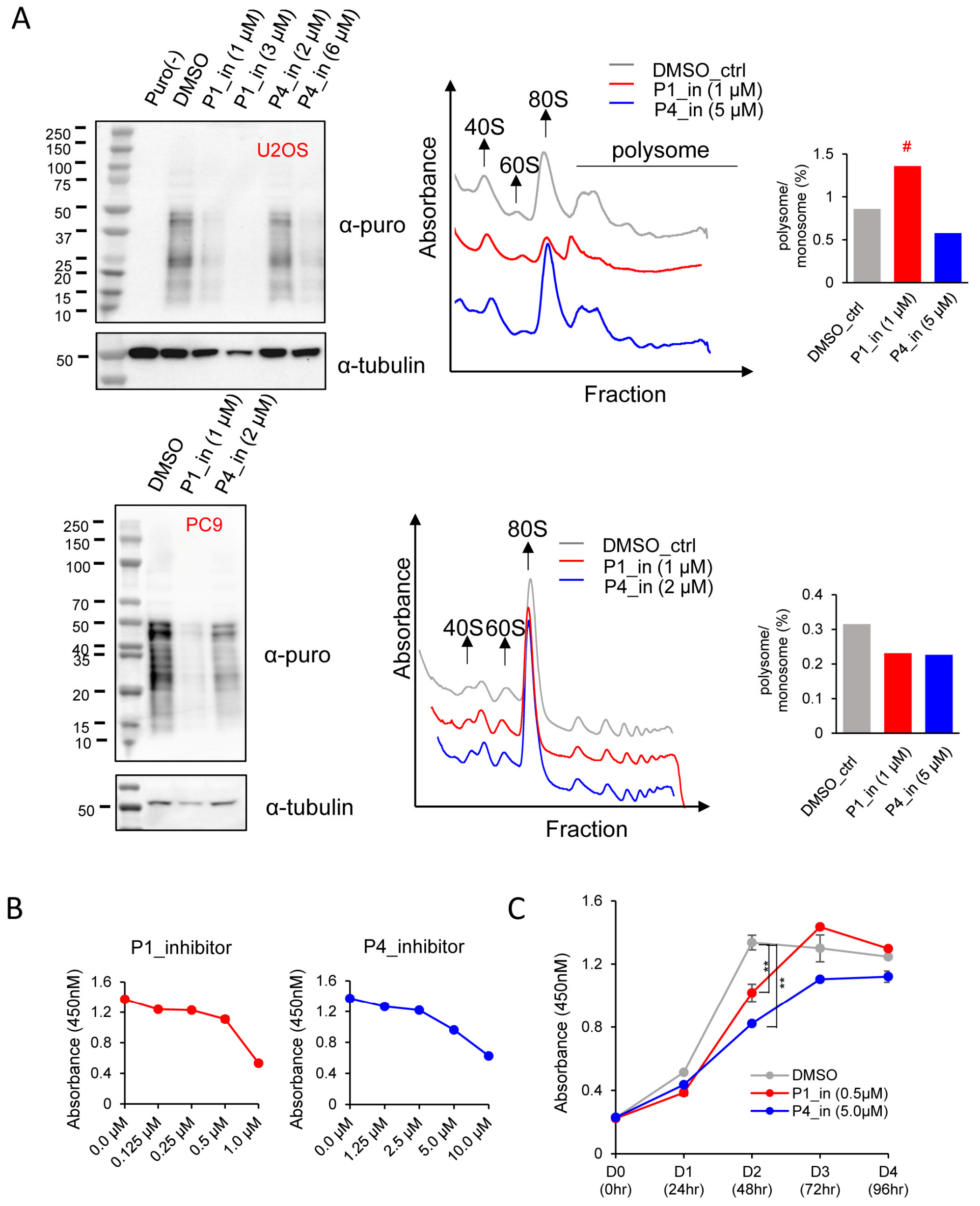


**Figure S7 (related to Figure 6). PRMT inhibition can lead to translation deficiency in other cell types and inhibition of cell proliferation.**

(A) U2OS (upper) and PC9 cells (lower) were treated with different inhibitors for 24h as indicated, and puromycin was added 30 min before cell harvest. Left, the incorporations of puromycin which indicates translation efficiency were detected by western blot using anti-puromycin antibody. Right, the effects of PRMT inhibitors on ribosome fractions were demonstrated by polysome profiling. The amount of mRNA bound in monosome or polysome were quantified with peak area. # showed that the ratio of monosome vs. polysome increased due to the reducing of monosome amount in this sample. HEK293T cells were treated with P1 or P4 inhibitor for indicated concentrations (B, 48hr) or time (C). Cell proliferation started at 4000 cell/well in 96-well plates and were detected with 10μl CCK8 for 2-hr incubation. Two biological replicates were performed and ** indicates the *p*-value is less than 0.01.

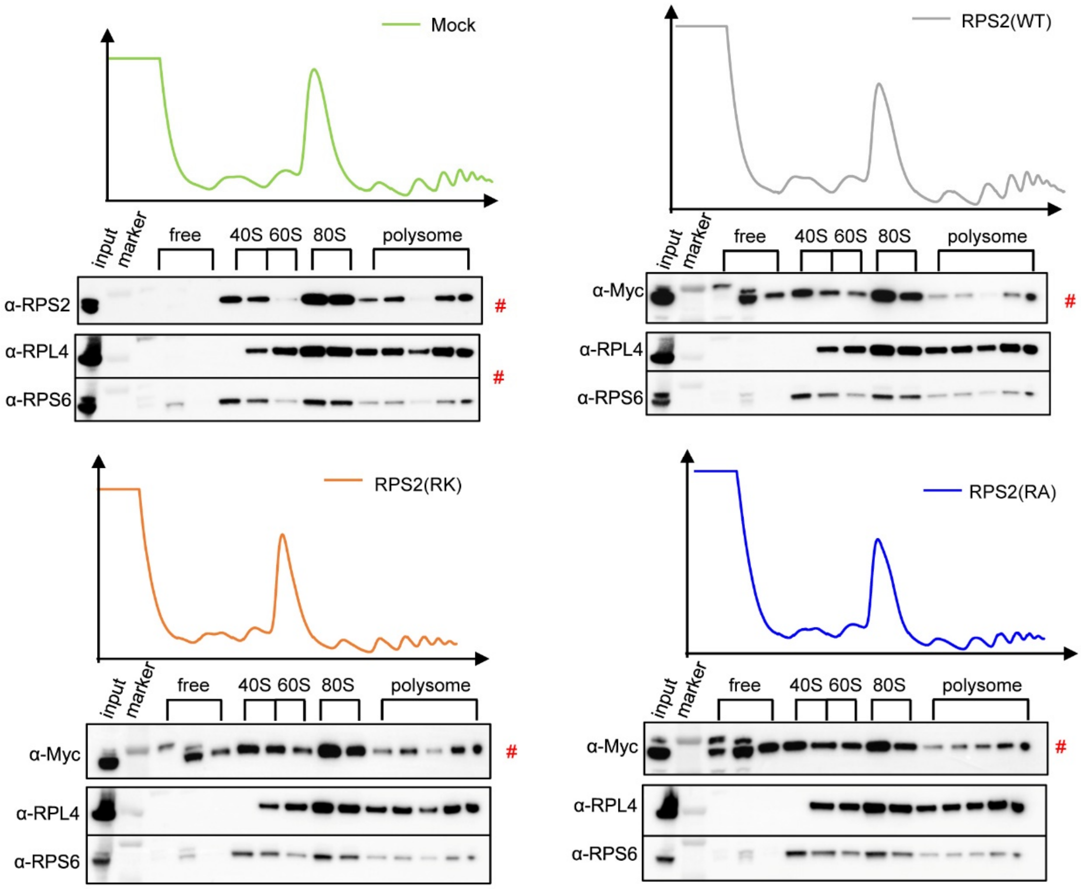


**Figure S8 (related to Figure 6). Arginine methylation of RPS2 affects ribosome assembly.**

Myc-RPS2(WT), Myc-RPS2(6RK) and Myc-RPS2(6RA) were transfected into HEK293T cells for 48hr, followed by polysome profiling. Fractions were collected and used for western blotting with anti-Myc, anti-RPS2, anti-RPS6 and anti-RPL4 antibodies as indicated (RPS6 and RPL4 were applied as profiling marker). Red # indicated what were selected to be shown in Fig. 6D
